# Spontaneous emergence and evolution of neuronal sequences in recurrent networks

**DOI:** 10.1101/2024.09.27.615499

**Authors:** Shuai Shao, Juan Luis Riquelme, Julijana Gjorgjieva

## Abstract

Repeating sequences of neural activity exist across brain regions of different animals are thought to underlie diverse computations. However, their emergence and evolution during ongoing synaptic plasticity remain unclear. To mechanistically understand this process, we investigated the interaction of biologically-inspired activity-dependent synaptic plasticity rules in models of recurrent circuits to produce connectivity structures that support neuronal sequences. Under unstructured inputs, our recurrent networks developed strong unidirectional connections, resulting in spontaneous repeating sequences of spikes. During ongoing plasticity these sequences repeated despite turnover of individual synaptic connections, a process reminiscent of synaptic drift. The turnover process occurred over different timescales, with sequence-promoting connectivity types and motif structures being strengthened while others weakened, leading to sequences with different degrees of volatility. Structured inputs could reinforce or retrain the resulting connectivity structures underlying sequences, enabling stable yet flexible encoding of inputs. Our model unveils the interplay between synaptic plasticity and sequential activity in recurrent networks, providing insights into how the brain might implement reliable and flexible computations.

## Introduction

Repeating sequences of neural activity have been observed across many species and brain regions, including in the navigational, sensory, and motor areas of mammals, reptiles, and birds, and are believed to underlie many computations. A striking example of repeating sequences is in the motor area of songbirds [1]. Zebra finches repeat their courtship song thousands of times with a perfectly matching sequential representation in their brain [2]. In the hippocampus of rats, sequences of spikes encode place trajectories as the animal moves, and they are replayed during rest [3, 4]. Repeating sequences of spikes are also common in the sensory cortices of rats, humans, mice, and turtles [5–8]. Yet, whether and how these sequences emerge and change over time to flexibly implement these computations remains poorly understood.

Sequences may repeat but are also subject to change. Growing experimental evidence shows that representations in multiple brain areas are often subject to “drift” [9–14] where one neuronal pattern of activations is slowly replaced by another. In the hippocampus, after the mapping of space is established [15–17] with the formation of place fields by training animals in the same environment, spontaneous changes in the encoding, a phenomenon known as “remapping”, redefine existing sequences [18, 19]. In the birdsong system, sequences are first learned and heavily modified by the juvenile bird as it develops its own version before the song crystallizes [20]. Even as an adult, the song can be modified under acute manipulations of their auditory system, indicative of the ongoing plasticity to which it is exposed [21, 22]. These sequences may be the result of a composition of smaller sequential units that can be replayed independently during sleep [23]. Even though the zebra finch is a specialist of one song, it still produces other non-courtship vocalizations in adulthood using the same motor areas [24–26]. Related species such as budgerigars, bengalese finches and canaries present more variable songs and even life-long learning [27–29]. Taken together, evidence suggests that the neuronal substrate underlying sequences may be more plastic than typically assumed.

The specific processes driving these changes in neuronal sequences are unknown, but likely relate to activity-dependent synaptic plasticity due to sensory experience [30, 31] or prior developmental spontaneous activity [32–34]. Indeed, many plasticity mechanisms have been studied, theoretically and experimentally, and are considered the main actors of organization of recurrent networks of neurons [35–45]. Experimental and theoretical evidence has put forward different forms of spike-timing-dependent plasticity (STDP) as the underlying plasticity mechanism which describes synaptic change based on the order and timing of pairs and triplets of preand postsynaptic spikes [36, 38, 42, 43, 45–48]. The synaptic plasticity induced by STDP can have different effects on recurrent network connectivity. For example, Hebbian forms of symmetric pair-based STDP, shifted asymmetric STDP, or rate-dependent triplet STDP have been shown to promote the formation of assemblies [42, 44, 49–52], groups of neurons sharing strong connections that exhibit coordinated firing [50, 53]. In contrast, anti-symmetric Hebbian STDP has been shown to promote the formation of “synfire chains”, a form of idealized sequential structures [37, 39]. Asymmetric Hebbian rules based on firing rates have also been discovered using meta-learning approaches capable of organizing and maintaining sequence-generating dynamics specifically applied to the HVC of zebra finches [54]. What specific synaptic plasticity rules promote structures of connectivity that support repeatable yet flexible neuronal sequences of cortical spikes is still unknown. Studying synaptic plasticity is challenging, especially in recurrent circuits, since activity and connectivity influence each other. Previous theoretical work has suggested that sequences provide stable output and robustly encode memories despite ongoing plasticity which modifies synaptic weights [55, 56]. However, if, as described above, neuronal representations are subject to constant change, driven through either internal dynamics or external stimuli, we lack an understanding of how this change affects the repeatability of spike sequences and their underlying structures.

Here, we asked how spike timing-dependent synaptic plasticity may produce the connectivity underlying repeatable sequences of spikes, and how this very same mechanism alters them. To mechanistically dissect this question, we built a recurrent, plastic, spiking network model that produced spiking sequences. Driven by unstructured activity, the network spontaneously developed connectivity structures through plasticity that led to the appearance of spiking sequences. These same plastic processes also created volatile dynamics of synaptic turnover with different timescales depending on synapse type, strength, and connectivity motifs. Exposing the network to structured inputs revealed that the same plasticity mechanisms could retrain the connectivity structures underlying sequences, retaining flexibility to encoding novel inputs. Overall, our work addresses how the interaction between spike timing-based plasticity rules and activity leads to the stability and evolution of spiking neuronal sequences which underlie diverse neuronal representations in neural networks.

## Results

### Plasticity drives strong synapses that produce repeating sequences

Previous work has shown that long-tailed distributions of synaptic strengths are sufficient to generate reliable but flexible spiking sequences reminiscent of those found in the cortex [57]. Thus, we first focused on identifying plasticity rules that produced such distributions in random networks and verified that they produced spiking sequences in recurrent neural networks with randomly connected excitatory (E) and inhibitory (I) adaptive exponential integrate-and-fire neurons [58] (Fig. 1A). We modeled spontaneous activity as uncorrelated Poisson inputs independently sampled for every neuron. E-to-E and I-to-E connections experienced synaptic plasticity according to spike-timing-dependent plasticity rules (eSTDP and iSTDP respectively, Fig. S1A-B) [37, 38, 46]. To prevent any small fraction of neurons from dominating the entire network, we applied weight normalization [59, 60] to E-to-E and I-to-E connections from both the presynaptic and postsynaptic sides to regulate the total input and output synaptic weights for each neuron, respectively (Fig. S1C). In addition, to further avoid dominance by individual neurons and to maintain a low overall firing rate comparable to that observed in the cortex during sequences [8], excitatory neurons were also regulated by intrinsic synaptic plasticity [61, 62]. This mechanism adjusted their firing thresholds based on their activity, thus stabilizing their firing rates over time (Fig. S1D).

**Figure 1.**
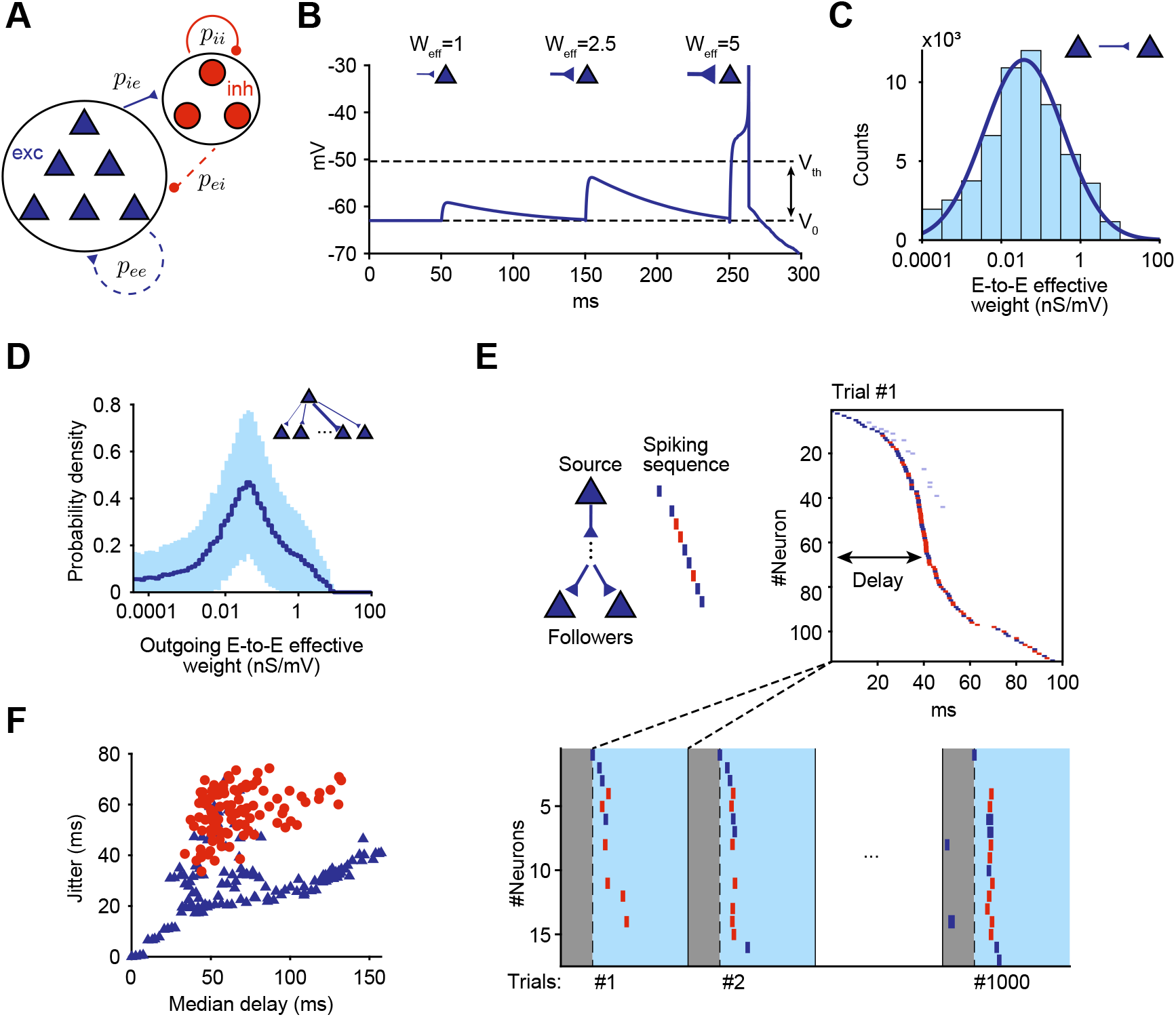
A plastic recurrent network can generate sequences under unstructured inputs. **A**. Schematic of the model network with 1,200 excitatory and 240 inhibitory AdEx neurons, where E-to-E and I-to-E connections are plastic. **B**. Example EPSPs initiated by connections with different *W*_eff_ on an isolated neuron. The effective weight of a connection (*W*_eff_) depends on the synaptic weight (*W* ), the postsynaptic resting potential (*V*_0_), and the firing threshold (*V*_*th*_). **C**. The network develops a lognormal E-to-E weight distribution in the steady-state. **D**. Different presynaptic neurons have similarly-distributed output connections. Solid line indicates the average and shade represents the standard deviation. Pooled over all E neurons in the network (*n* = 1, 200). **E**. Top: When a randomly chosen “source” neuron is forced to spike, its “followers” activate reliably and sequentially. Neurons are sorted by their spike time in example trial. Blue and red ticks indicate excitatory and inhibitory followers. Bottom: the source neuron was forced to spike in 1,000 consecutive trials and increases in firing rate were calculated to identify reliable followers. **F**. The jitter of the followers is positively correlated with the median delay. Same followers as E.

Since intrinsic plasticity adjusts the firing thresholds, we studied synaptic strengths after scaling them by the distance between the postsynaptic neuron’s threshold and resting potentials. We refer to this value as *effective* weight (*W*_eff_ = *W/*(*V*_*th*_ − *V*_0_)), which provides a good indicator of the likelihood that a presynaptic activation would trigger a postsynaptic spike (Fig. 1B). The steady state of the plastic network subjected to uncorrelated inputs resulted in an E-to-E effective weight distribution that was skewed, with a long tail well described by a lognormal distribution (*R*^2^ = 0.94, Fig. 1C). We found that the rare strong connections in the tail of the distribution were distributed among the entire excitatory population, with most excitatory neurons having at least one such output connection (Fig. 1D, S2A).

Following previous work [8,57], we tested the generation of sequences by randomly selecting an excitatory neuron (source neuron) in the model network, injecting input into it to produce 1,000 spikes at regular intervals which defined a trial (Fig. 1E) and measuring the response of all other neurons in the network. We quantified the resulting firing rate modulation as the difference between each neuron’s firing rate after and before the source neuron spikes. Consistent with prior work [8, 57], we defined as “followers” those neurons that exhibited a significantly high firing rate modulation when compared to a null model with no injected spikes (Methods). Out of 155 tested source neurons in a model network, 135 (87.1%) of them had at least one excitatory follower. The median number of excitatory followers (excluding 0) was 55, and source neurons could have more than 120 excitatory followers (Fig. S2B).

Source spikes caused the activation of followers in the same sequence across trials (Fig. 1E, S3, and S4, [8]). Followers could activate after the source spike with a delay up to 150 ms, much longer than the timescales of neuronal dynamics. Different subsets of all followers fired in every single trial (55% excitatory and 74% inhibitory followers per trial on average, S2C). We thus defined the responding probability of a follower as the ratio out of all trials in which it fired at least once within 300 ms from the source spike. Across all followers, the median delay and the jitter increased together, while both of them decreased with increasing responding probability (Fig. 1F, Fig. S5).

Surprisingly, although the model lacked E-to-I plasticity and thus could not produce strong connections onto inhibitory neurons, it did produce inhibitory followers (median: 33, 45.8% of source neurons) (Fig. S2). Compared to excitatory followers with the same responding probability or median delay, inhibitory followers had a larger jitter (Fig. 1F, S4 and S5C). We found that the activation of inhibitory followers results in feedback inhibition onto the source neuron (Fig. S6), possibly explaining post-spike hyperpolarization observed experimentally [8].

In summary, plasticity in our network generated connectivity that promotes the appearance of sequences even under random inputs, consistent with experimental evidence in cortex [8]. Single spikes in most excitatory neurons reliably trigger sequences of spikes in the rest of the network that repeat from trial to trial.

### Strong excitatory connections resist turnover under uncorrelated inputs

Although our model network converged to a steady state in terms of the distribution of effective weights, individual connections could still change due to ongoing plasticity (distribution and single example connections tracked in Fig. 2A). Since spike sequences in our networks were mainly driven by strong connections, we wondered if these changes could limit the capacity of the network to produce repeatable sequences over long timescales.

**Figure 2.**
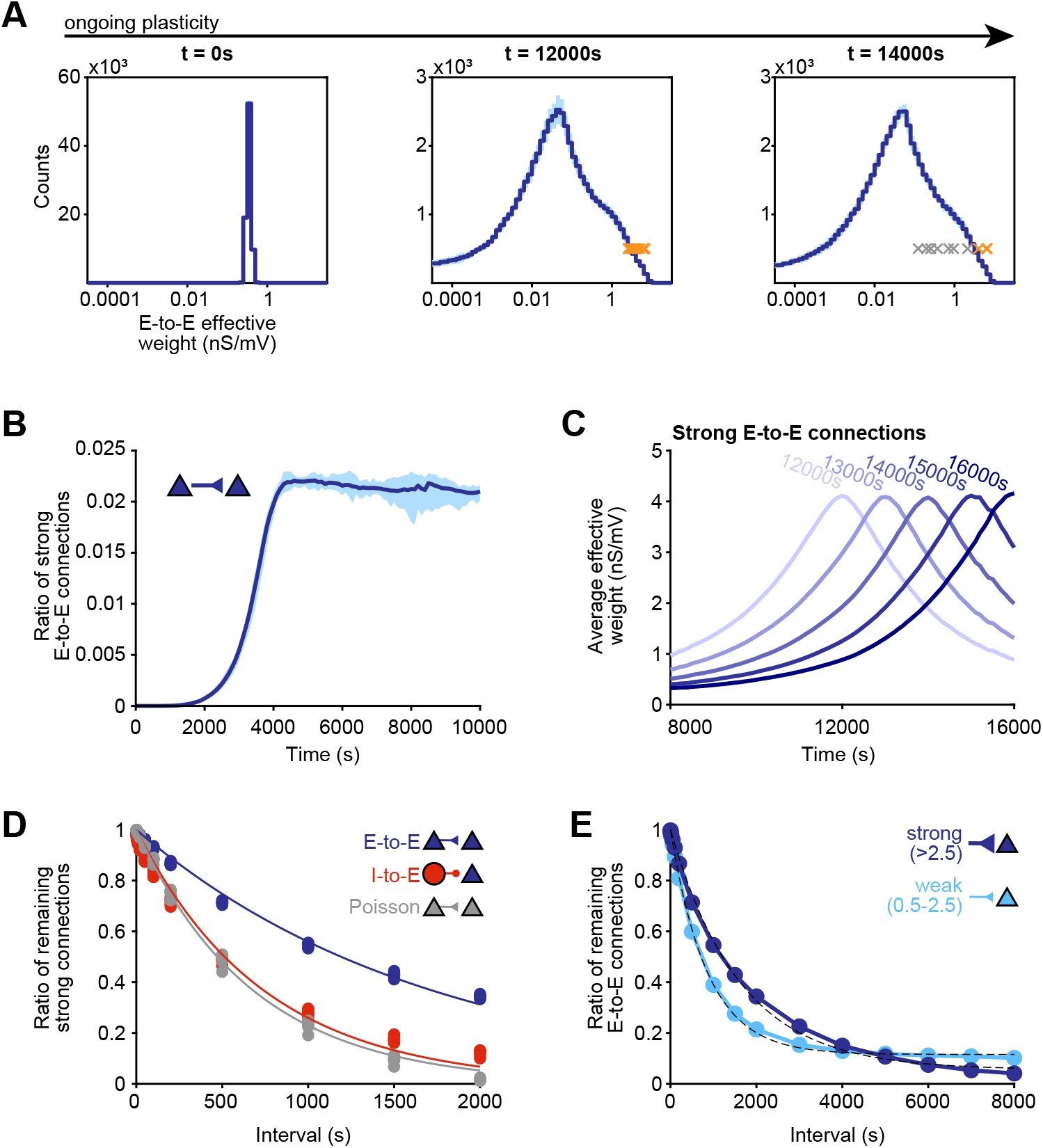
Turnover of strong connections in the plastic network. **A**. Evolution of weight distribution (line and shading; mean ±0020std) and example single connections (crosses) at multiple time points. Colored crosses indicate strong connections (*W*_eff_ *>* 2.5). The distribution reaches a steady state, but single connections continue to fluctuate in strength. **B**. Ratio of strong E-to-E connections over time. **C**. Average strength of multiple groups of E-to-E connections randomly picked from the tail of the distribution at different time points (colored labels), after the weight distribution has reached a steady state. **D**. Decay of strong E-to-E (blue) and I-to-E connections (red). Dots indicate the ratio of strong connections that remain strong after a given interval (abscissa). Lines indicate exponential fits. Gray represents E-to-E connections when neurons are forced to fire Poisson spike trains. *τ*_*ee*_ = 1, 719 s, *τ*_*ie*_ = 741 s, *τ*_*poisson*_ = 679 s, *n* = 10 networks. **E**. Decay of strong E-to-E (*W*_eff_ *>* 2.5, same as the average of blue points in **D**, up to 2,000 s) and weak E-to-E connections (0.5 *< W*_eff_ ≤2.5, *n* = 10 networks). Dashed lines indicate exponential fits with baseline.

We first defined the ratio of strong E-to-E connections to the expected number of E-to-E connections (*p*_*ee*_*N*_*e*_(*N*_*e*_ −1), Fig. 2B). Once the steady state was reached, our network could always produce sequences as a consequence of a stable number of strong connections distributed over the network independent of their exact identity. When we tracked small groups of E-to-E connections that lived in the tail of the distribution at a given time point (*W*_eff_ *>* 2.5, Fig. S2A), we found that they had been weak in the past and would become weak again in the future (Fig. 2C). This group would then be replaced by a new group of connections preserving the tail of the distribution. Strong I-to-E connections followed the same dynamics (Fig. S7A). Indeed, under uncorrelated inputs, the probability that a connection remained strong after a given period decayed exponentially (Fig. 2D, blue).

To quantify the rate of this turnover of strong connections, we first simulated a baseline network under the same plasticity rules where all of the neurons generated independent Poisson spikes with the same rate as our network simulations (Fig. 2D, gray). We found that the turnover of strong connections was much slower than this random baseline. This slowed turnover is likely the product of the synaptic potentiation resulting from the anti-symmetric eSTDP rule combined with positive spiking correlations between pre- and postsynaptic units. Indeed, I-to-E connections, subject to symmetric iSTDP and negative correlations resulting from inhibition, experienced similar turnover rates to the baseline network (Fig. 2D, red).

Strong E-to-E connections did not just experience slower turnover compared to I-to-E, but also when compared to weak E-to-E connections in the body of the distribution. We quantified the turnover of weak connections (*W*_eff_ ∈ (0.5, 2.5]) as the ratio of connections that remained above their lower bound after some elapsed time. The ratio first decreased and then saturated (Fig. 2E and S7B), and could be described with an exponential function and a baseline, i.e., 1 − *a* [1 − exp(−*t/τ* )]. The time constant for the turnover for weak connections (*τ* = 856 s) was of the same scale as in the baseline network and I-to-E fit (Fig. 2D, gray and red) and again much shorter than that of strong connections (*W*_eff_ *>* 2.5; *τ* = 1, 605 s). Interestingly, I-to-E connections showed the opposite trend: strong connections turned over slightly faster (*W*_eff_ *>* 2.5; *τ* = 599 s) than weak ones (*W*_eff_ ∈ (0.5, 2.5]; *τ* = 781 s) (Fig. S7B).

In summary, under uncorrelated inputs and ongoing plasticity, our networks displayed stability at the global level of synaptic strength distributions, but volatility of individual connections. While all connections experienced turnover in their strength, the strongest E-to-E connections displayed slower decay than the rest of the network. These strong connections thus provide a backbone underlying the generation of spiking sequences.

### Plasticity promotes connectivity motifs involved in sequences

Studies have shown that certain connectivity motifs are more likely to appear or be activated in recurrent cortical networks [57, 63–66]. Given the slow turnover and relevance of strong connections in the generation of spike sequences, we next investigated how plasticity structures the connectivity motifs of strong connections in particular.

We focused on four 3-neuron motif types composed of strong E-to-E connections: linear chains, divergence motifs, convergence motifs and fan-in/out motifs (*W*_eff_ *>* 2.5, Fig. 3A). While another 3-neuron motif is also possible (a cycle motif, where all three neurons are connected in a recurrent loop [65, 67]) due to the asymmetry of the eSTDP rule, it was almost never observed in our network (similar to [67]), with an average of 0.08 motifs per network instance. Following previous work, we defined motif frequency as the ratio of the observed number of motifs to the expected number for each motif type (see Methods for details on each motif) [68–70]. During ongoing plasticity in our network, we initially observed a sudden increase in the frequency of motifs for all motif types, which then settled at a steady state (Fig. 3B). This initial increase mimicked that observed for single strong connections (Fig. 2B), but was not merely a consequence of it. Compared to the motif frequency in a random network, given the same probability of strong unidirectional connections, we found that the networks in steady-state had over-represented “fan-in/out” motifs but under-represented all other motif types (Fig. 3C).

**Figure 3.**
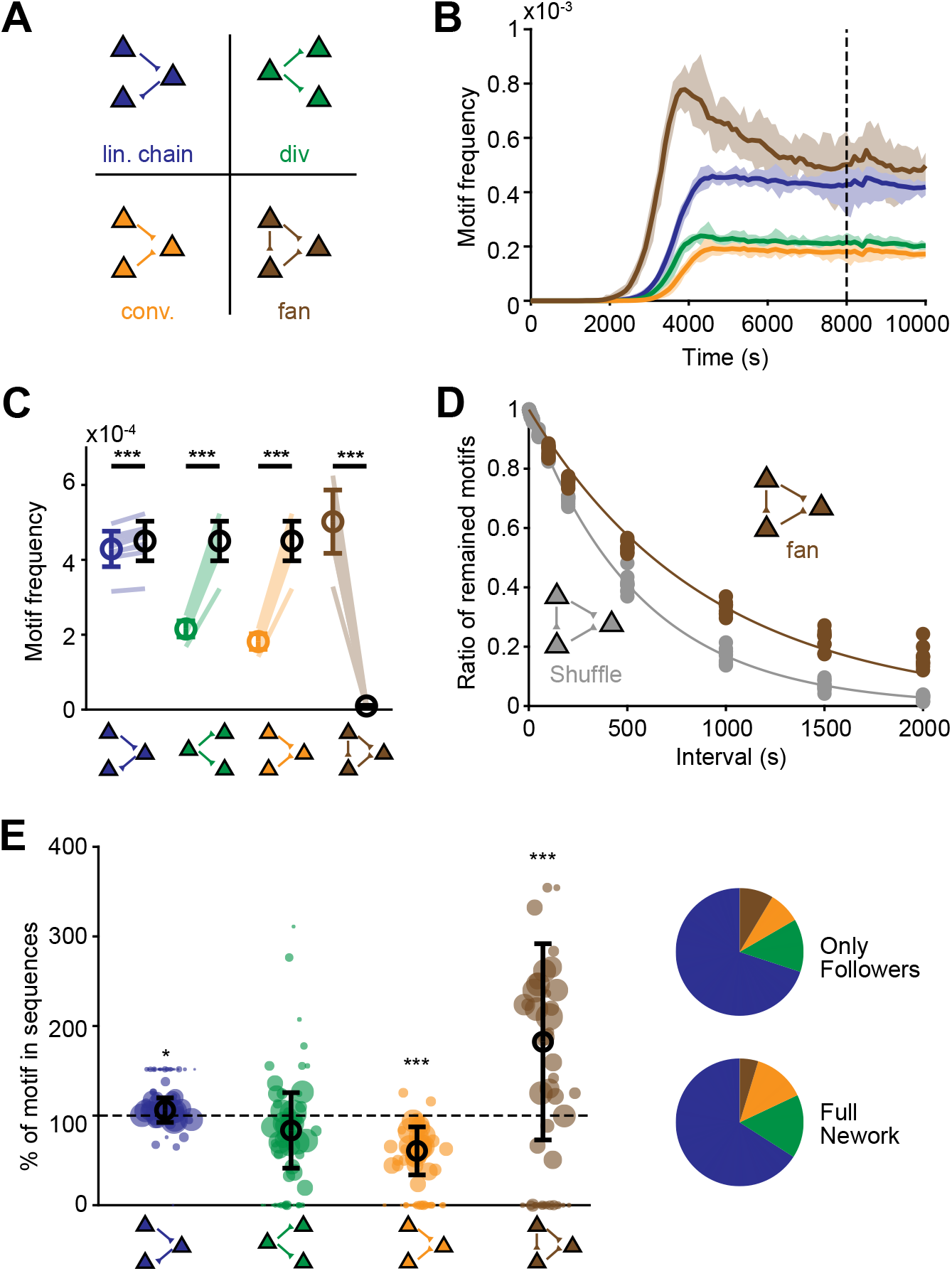
Motif analysis of strong connections. **A**. Definitions of the four triplet motifs: linear chains, divergence motifs, convergence motifs, and fan-in/out motifs. **B**. Frequency of triplet motifs of strong connections over time, relative to the number of all possible sites to form a motif. The dashed line corresponds to the data points in **C. C**. Motif frequency in our model network (colored) compared to the estimated motif frequency in a random network with the same ratio of strong E-to-E connections (black) (*n* = 10 networks; mean ± std). ***: *P <* 0.001, paired sample t-test. **D**. Decay of the fan-in/out motif (colored) compared to a null model (gray) with shuffled connections. Lines indicate exponential fits (*τ*_fan_ = 907 s, *τ*_shuffle_ = 562 s). **E**. Percentage of motifs among followers relative to the percentage among all neurons in the network (*n* = 74 sequences; mean ± std). Pie charts: percentages among followers (linear chains: 69.95%; divergence: 13.42%; convergence: 8.05%; fan-in/out: 8.58%, summing up to 100%) and in the full network (linear chains: 65.94%; divergence: 16.08%; convergence: 13.28%; fan-in/out: 4.70%, summing up to 100%). *: *P <* 0.05, ***: *P <* 0.001, single sample t-test.

We hypothesized that the presence of convergence and divergence motifs was limited by the competition among multiple connections received by a single neuron that results from weight normalization. Additionally, by regulating the firing rate of all neurons, intrinsic plasticity would also discourage the clustering of strong connections on single neurons required for these motifs. To verify these hypotheses, we disabled the presynaptic weight normalization and intrinsic plasticity on E neurons in new simulations. We found that the motif frequency of all types in these alternative models were above random levels (Fig. S8A), but the ratio of strong connections was lower (2.1 × 10^−3^ vs. 0.021). This alternative model contained a small number (4%) of “hub” neurons that receive from, or project to, other neurons most of the strong connections of the network. These hub neurons also fired much more frequently than average, exerting a large impact on the network dynamics (Fig. S8B, [71–73]). In these models, ∼5% of the excitatory neurons (mainly the hub neurons) had followers and their activation was very short lived (*<*20ms), closer to synchronous firing than to sequential propagation (Fig. S8C-D). Therefore, the plasticity rules that promote the homogeneous distribution of strong connections required for the generation of spiking sequences simultaneously limit the presence of divergent and convergent motifs. If this argument were true, then fan-in/out motifs, which also require clustering of strong connections on single neurons and can be transformed into any of the other three motifs by removing just one strong connection, should also be limited by the plastic mechanisms described above. Given that single strong connections are subject to turnover (Fig. 2B), it is thus surprising that fan-in/out motifs were so over-represented.

A hypothesis for the over-representation of fan-in/out motifs is self-reinforcement due to ongoing synaptic plasticity. To test this, we explored the turnover rate of the fan-in/out motif relative to single strong connections. We established a baseline where we exchanged a random selection of strong connections with weak ones to achieve the same turnover rate of individual strong E-to-E connections as before (Fig. 2D). The decay time constant of the shuffled fan-in/out motifs was approximately 1/3 of the time constant of strong unidirectional connections under normal turnover (i.e., 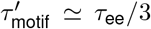) (Fig. 2D). In contrast, fan-in/out motifs turned over more slowly than this baseline (Fig. 3D). Therefore, the slow turnover of the fan-in/out motif could not be ascribed exclusively to the slow turnover of single strong connections, but rather indicates the existence of higher-order self-reinforcing effects, likely driven through the effect of eSTDP. When we performed equivalent analysis for the other motif types, we observed faster turnover compared to baseline (Fig. S9), showing that these motifs are not only less common but also highly volatile.

We then evaluated whether these motifs supported the generation of spiking sequences specifically. We considered only the E-to-E connections among the source neuron and its most reliable followers (responding probability ≥ 0.8). We calculated the relative percentage of each motif type (Fig. 3E, top pie chart), then normalized the motif percentages in the sequences by the motif percentages for the full network (Fig. 3E, bottom pie chart). We found that the fan-in/out motifs appeared even more frequently among followers than in the full network, where they are already over-represented (Fig. 3E, dots).

Taken together, our results show that plasticity rules produce specific structures of connectivity that support the reliable generation of spiking sequences. Fan-in/out motifs of strong connections self-reinforce, resulting in a slow turnover of the motif and their over-representation across the entire network and among followers. Convergence and divergence motifs on the other hand are suppressed through weight normalization and intrinsic plasticity, which promotes an homogeneous distribution of strong connections within the network.

### Turnover of strong connections causes turnover in sequence composition

While the observed turnover of strong E-to-E connections and fan-in/out motifs could cause the destruction of any particular sequence, the distribution of synaptic strengths remained stable, suggesting a constant replacement by new sequences. Indeed, sequences initiated by the same source neuron at different time points exhibited different follower composition, where some followers remained, some were lost, and new ones appeared (Fig. 4A). We compared new and lost followers within 100 s intervals and averaged over 50 sequences (Fig. 4B). The net increase in the number of followers (new - lost) in a 100 s interval was −0.1 ± 30.6, indicating that the size of the sequences remained stable over time.

**Figure 4.**
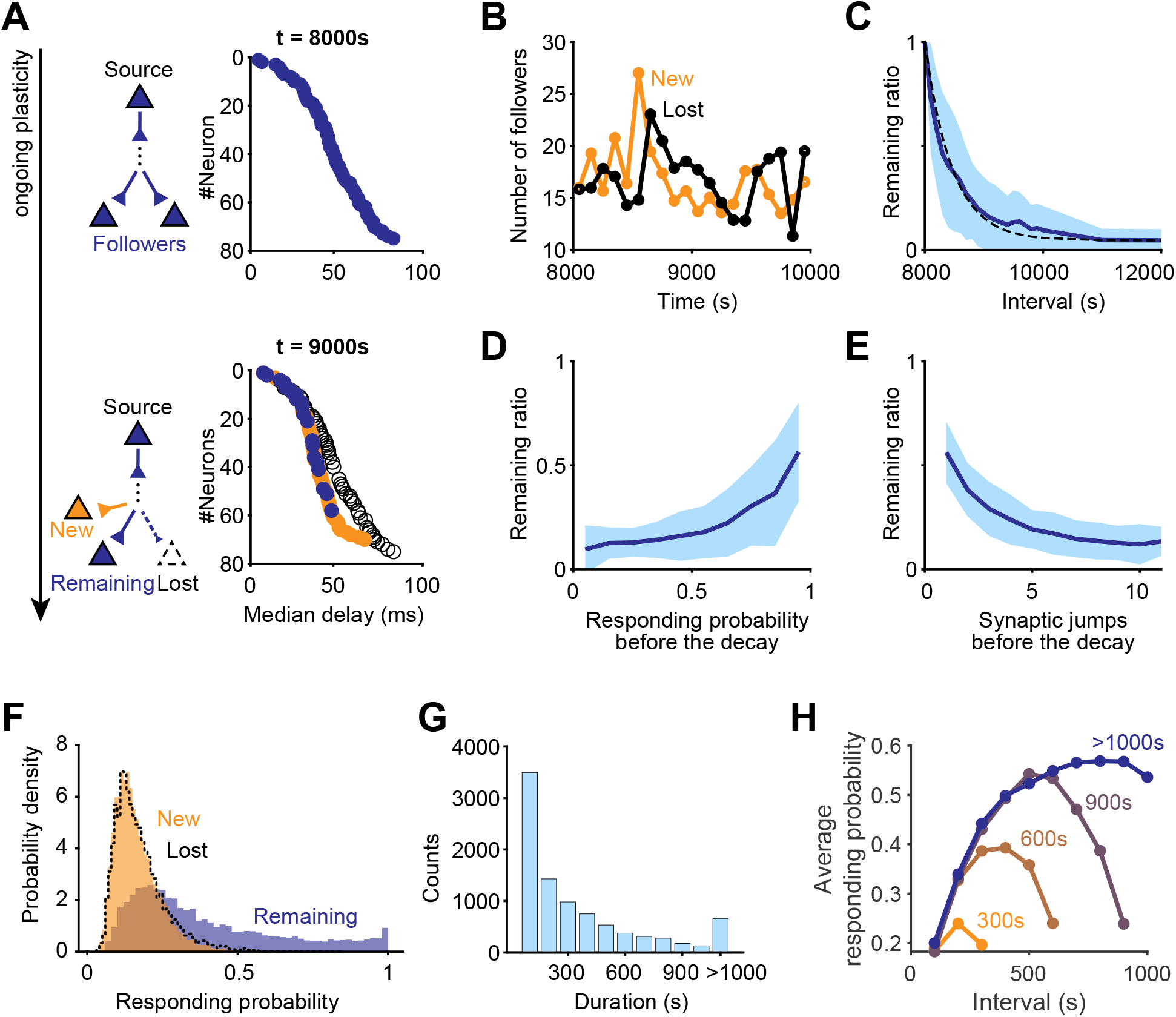
Turnover of followers and sequences. **A**. Excitatory followers of an example source neuron at two time points. Blue circles indicate common followers at both time points (Remaining). Hollow circles indicate cells classifying as followers earlier but not later (Lost). Orange circles indicate those classifying as followers later but not earlier (New). **B**. Mean number of new and lost followers over 50 sequences in 100 s intervals. **C**. Decay of the probability that a follower continues to classify as a follower after a given interval (pooled over 2,931 followers from 50 sequences, mean ± std). Dashed line indicates exponential fit with baseline. **D**. Ratio of remaining followers after 1,000 s as a function of their responding probability (pooled over 24,311 followers from 50 sequences at 8 time points, mean ± std). **E**. Same as **D** but as a function of number of synaptic jumps from the source. **F**. Distributions of the responding probability for new, remaining, and lost followers in 100 s intervals. **G**. Distribution of the followers’ lifetime. Pooled from followers detected every 100 s from 50 different sequences. **H**. Mean responding probability for different groups of followers, grouped by by their lifetime. Pooled over 2,338 followers from 50 sequences.

To evaluate the rate of follower turnover, we first pooled 2,931 followers from 50 randomly selected source neurons over 10 model networks which reached steady state after training with the same plasticity rules and parameters. We then re-evaluated if these neurons continued to classify as followers at regular intervals during ongoing plasticity. We found that the ratio of initial followers that remain followers dropped exponentially (Fig. 4C).

Follower turnover was much faster than that of single strong connections (Fig. 2D). To characterize the differences in turnover rates among followers, we analyzed followers based on their responding probabilities and delays. We found that the ratio of remaining followers was the highest for followers with the highest responding probability and shortest delay (Fig. 4D, Fig. S10A). Considering that strong E-to-E connections are the main conduit of follower activation (Fig. 1, [57]), we calculated the length of the shortest path connecting each follower to its source using only strong E-to-E connections (which we call “synaptic jumps”). The length of this path negatively correlated with the follower’s responding probability and positively correlated with its median delay (Fig. S10B-C). Consistent with these correlations, the remaining ratio of of followers also decreased with an increasing number of synaptic jumps (Fig. 4E). In summary, followers synaptically closest to the source neuron respond most reliably and are the most robust to turnover.

We next asked how long followers last and whether their properties evolve with ongoing plasticity. We divided a period (2,000 s) after reaching steady state into small intervals (100 s) and detected new, lost and remaining followers. In each of these intervals, new followers had lower responding probabilities than those that remained from the previous interval, as well as longer delays, and more synaptic jumps from the source neuron. The distribution of the responding probability of lost followers in every interval overlapped with that of new followers (Fig. 4F), confirming that the overall number of followers in the network remained stable (Fig. 2A). When pooling new followers from all intervals (*n* = 7, 189), we observed a skewed distribution of lifetimes (Fig. 4G), with most neurons being classified as followers for only a short period. Interestingly, a small proportion (7.3%) of new followers lasted over 1,000 s, much longer than the time constant of the overall turnover of followers, *τ*_*seq*_ = 459 s (Fig. 4D), suggesting that the network may stabilize some new followers after they are recruited. The responding probability of followers followed a non-monotonic trajectory, first increasing and then decreasing with time, which was common to all followers, independent of their lifetime (Fig. 4H). Some followers displayed remarkably long lifetimes, remaining followers for as long as our simulations. These highly stable followers also showed peak responding probability.

In summary, in the presence of uncorrelated inputs and ongoing synaptic plasticity, our networks showed stable global properties and local fluctuations, not only for single synaptic connection and motif distributions, but also for sequence composition. Single sequences experienced the appearance of new followers and disappearance of old ones so that the overall extent of spiking sequences in the network remained constant. While many followers experienced short lifetimes with low responding probabilities, others could be stabilized in the network, becoming long-lasting and highly responsive followers.

### Spatially structured inputs reinforce strong connections and stabilize turnover

To investigate whether spiking sequences might also be generated in the presence of structured inputs, as is the case in sensory areas which predominantly receive and process sensory input, we next drove the networks with correlated inputs in the presence of the same types of synaptic plasticity. We applied a common training paradigm that is used to form assemblies, defined as strongly mutually connected groups of neurons that experience correlated activity [35,40,42,43,50,51,53,74,75]. We refer to these as ‘spatially’ correlated inputs, implemented as additional inputs to ongoing unstructured activity sequentially targeting groups of excitatory neurons (which became the assemblies) one group at a time, with a different group being targeted every 100 s (Fig. 5A).

**Figure 5.**
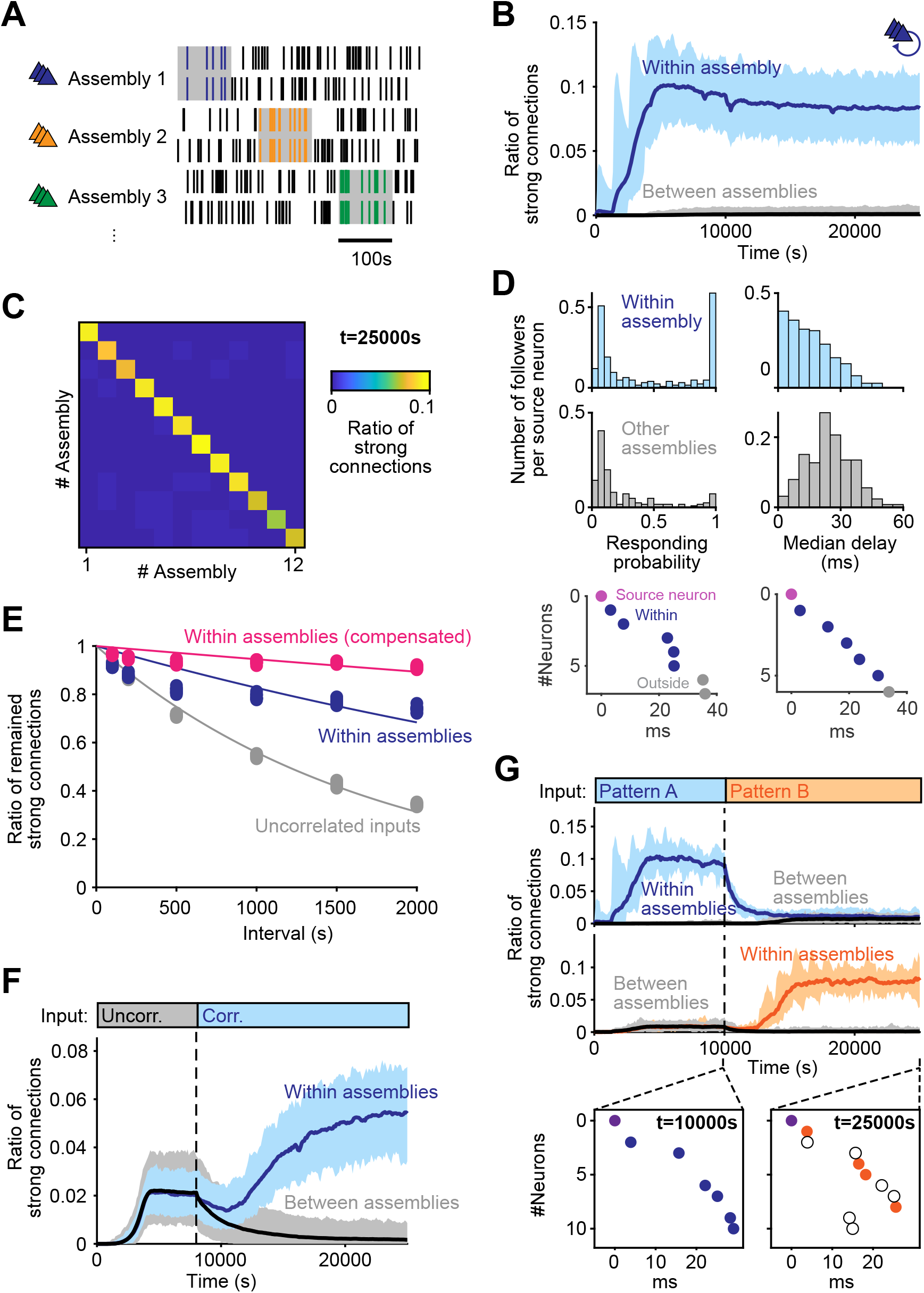
Structured inputs lead to structured connectivity and stable turnover. **A**. Schematic of training protocol with spatially structured inputs. Correlated inputs were sequentially presented to non-overlapping groups (which determined the assemblies) of excitatory neurons (color-coded) for 100 s (gray shading). Only one assembly received inputs at a time. Other excitatory and inhibitory neurons received uncorrelated inputs. The targeted group cycled throughout the simulation. **B**. Ratio of strong E-to-E connections where pre- and postsynaptic neurons are in the same assembly (within) or in different assemblies (between), (*n* = 10; mean ± std). **C**. Ratio of strong connections within and between assemblies at steady state (*t* = 25, 000 s in B). Most of the strong connections are formed within the correlated assemblies (diagonal). **D**. Top: Distribution of responding probability (left) and median delay (right) of followers in the same assembly with the source neuron (blue, *n* = 155) or in other assemblies (gray, *n* = 97). Bottom: Two example sequences triggered at two source neurons in different assemblies. Purple: source neurons. Blue: Followers within an assembly. Gray: Followers in other assemblies. **E**. Decay rate of strong E-E connections in the network with and without correlated inputs. Blue: E-to-E connections within assemblies. Gray: Decay in the network with uncorrelated inputs (same as Fig. 2D, blue). Pink: Same as blue but turnover of a connection was not counted if it was replaced by a new connection in the same assembly. Lines indicate exponential fits (*τ*_*within*_ = 5, 259 s, *τ*_*uncorr*_ = 1, 721 s, *τ*_*comp*_ = 17, 761 s). **F**. Ratio of strong connections, in networks first trained with uncorrelated inputs and then correlated inputs (*n* = 10; mean ± std). **G**. Same as **F** but networks were first trained with one correlated input pattern and then switched to a different correlated input pattern. Top: within and between connections relative to first pattern. Middle: same relative to second pattern. Bottom: Example sequences from the same source at two time points. Orange and hollow circles are new and lost followers relative to the first time point.

The network reached a stable long-tailed distribution of synaptic strengths (Fig. S11A-B). As a result of the spatially structured inputs, however, the strong E-to-E connections concentrated within single assemblies, with very low chance (9.1 × 10^−4^) of strong connections between different assemblies (Fig. 5B-C). Consequently, followers were most likely found within the same assembly (Fig. 5D). Pooling from 126 source neurons, we detected 252 followers in the same assembly as their source and 155 followers in any other of the remaining 11 assemblies. The percentage of same-assembly followers (61.9%) was much higher than what would be expected if followers were distributed uniformly in the entire network (9.1%). Furthermore, same-assembly followers had significantly higher responding probability than different-assembly followers (0.47 ± 0.40 vs. 0.25 ± 0.28) and shorter delays (15.7 ± 10.6 ms vs. 24.8 ± 10.8 ms). Hence, consistent with prior work on assemblies [40, 42, 43, 51, 74, 75], under spatially correlated input directed at different groups neurons, our networks also formed neural assemblies; yet, within each assembly, synaptic plasticity still generated strong connections. Moreover, the strong connections were concentrated within assemblies, reinforcing connectivity structures underlying sequences.

Spatially structured inputs also made strong connections more stable than those resulting from training under unstructured activity (as earlier in the study) (Fig. 5E, blue and gray). Furthermore, while turnover of single strong connections did occur, those decayed strong connections were quickly compensated by new ones in the same assembly, preserving the high density of strong connections within assemblies and resulting in very stable connectivity (Fig. 5E, pink).

We next asked if the stable strong connections concentrated within assemblies could be learned by a network that had been previously trained with unstructured activity (as earlier in the study), effectively retraining the original network structure. After training a network with unstructured inputs until it reached steady state (*t* = 8, 000 s), we again provided spatially correlated inputs to groups of neurons (Fig. 5F). We found that the network connectivity could indeed be retrained so that the strong connections between neurons underlying sequence generation reorganized from being broadly distributed across the entire network to being concentrated within the assemblies specified by the structured input. Correspondingly, the percentage of same-assembly followers rose from 8.3% to 42.9% (Fig. 5F).

Finally, we investigated the capacity of the network to adapt to a change in the inputs. After training a network with one pattern (A) of spatially correlated inputs until it reached steady state, we switched to a new, randomly chosen pattern (B) (Fig. 5G). The network restructured its connectivity with the new strong connections being concentrated within the assemblies specified by the new pattern B of spatially correlated inputs. Correspondingly, the percentage of same-assembly followers to pattern A dropped from 78.7% to 9.0%, while the percentage of same-assembly followers of pattern B rose from 5.5% to 68.8%. This redistribution of strong connections to the new assemblies was also reflected in the generated sequences. In summary, spatially structured inputs, providing correlated input to groups of neurons, could further shape network connectivity that leads to sequences. The strong connections underlying sequence generation concentrated between neurons receiving the correlated inputs and forming the assemblies, yielding reliable followers and slowing down turnover. Despite this enhanced stability, the network could be re-trained, remaining flexible to accommodate input changes.

### Temporally structured inputs define and stabilize cortical sequences

Cortical inputs may also contains temporal structure in addition to activating neurons in the same group. We thus explored the stability and flexibility of connectivity structures underlying sequences in networks driven by temporally correlated inputs during synaptic plasticity. These inputs were implemented as Poisson spikes targeting one assembly at a time and sequentially shifting to the next. Each assembly received inputs for only 10 ms—a timescale comparable to single-neuron dynamics in our model and much shorter than that used in the previous section (100 s). Differing from the previous section, an assembly received no input between activations, until all other assemblies had been activated and the sequentially shifting inputs returned to it (Fig. 6A). This resulted in the groups of neurons being sequentially targeted in the same quick succession during training, exposing neurons to strong temporal correlations. The network reached a stable long-tailed distribution of synaptic strengths (Fig. S11C-D). With these temporally structured inputs, strong connections emerged not only within assemblies (diagonal of the connectivity matrix), but also from one assembly to another (sub-diagonal, Fig. 6C). These next-assembly connections extended sequences over multiple assemblies (Fig. 6D).

**Figure 6.**
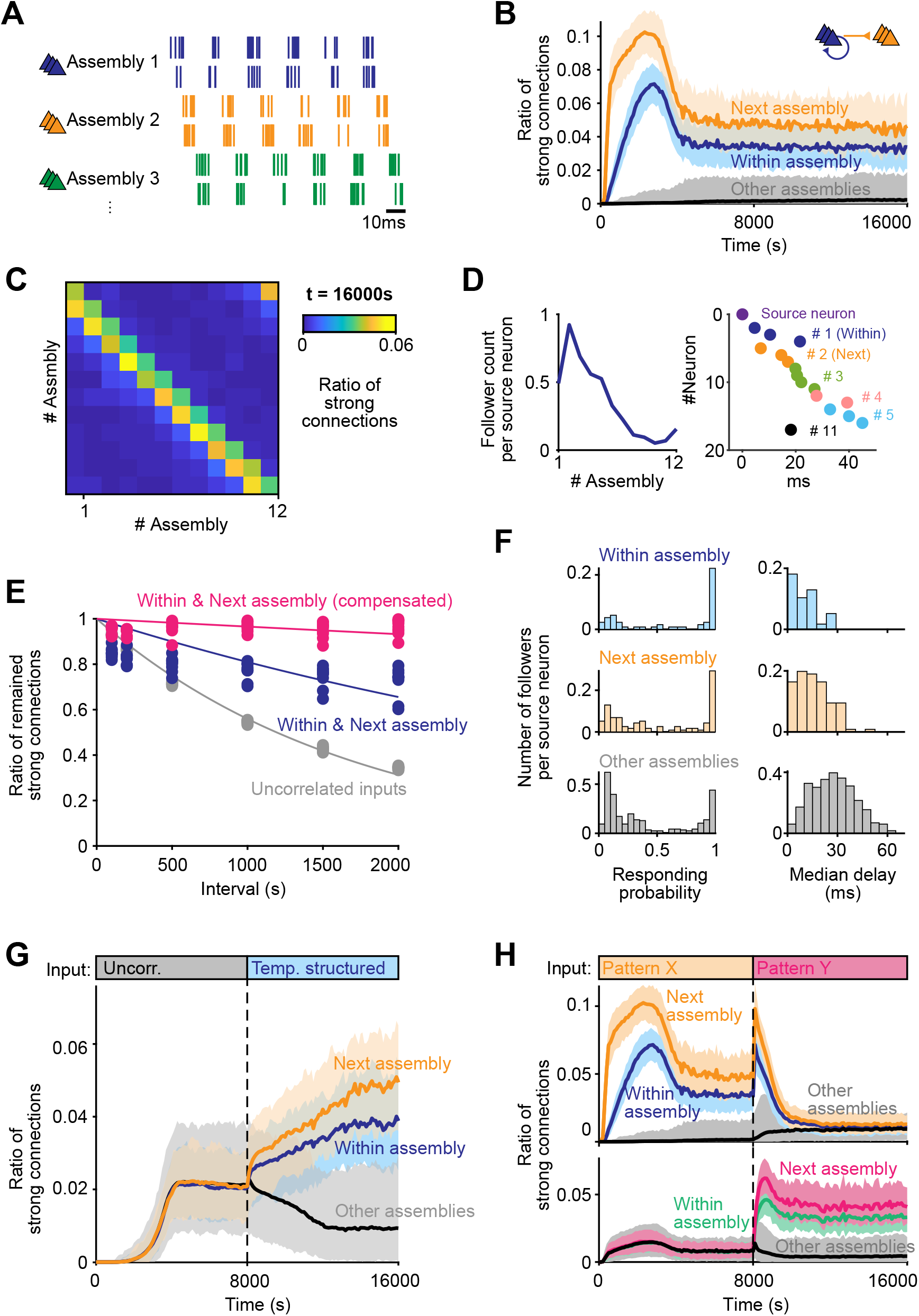
Temporally structure inputs lead to sequentially organized followers. **A**. Schematic of training protocol with temporally structured inputs. Correlated inputs were sequentially presented to non-overlapping groups (which determined the assemblies) of excitatory neurons (color-coded) for 10 ms (gray shading). Only one assembly received inputs at a time. Inhibitory neurons received uncorrelated inputs. The targeted group cycled throughout the simulation. **B**. Ratio of strong E-to-E connections within the same assembly (blue), from one assembly to the next assembly (orange), and all others (black) (*n* = 10; mean ± std). **C**. Ratio of strong connections within and between assemblies at steady state (*t* = 16, 000 s in B). **D**. Left: Average count of followers in different assemblies. Assembly number is relative to the source neuron, starting from 1. Note most followers fall in the immediate next assembly. Right: example sequence extending from assembly 1 to 5, with followers vertically sorted by the assembly number relative to the source neuron, as indicated by different colors and numbers. **E**. Decay rate of strong E-to-E connections. Blue: to a neuron in the same or next assembly. Pink: same as blue but turnover is counted only if the connection was not replaced by a new one. Black: Decay in the network with uncorrelated inputs (same as Fig. 2D). Lines indicate exponential fits (*τ*_*ee*_ = 4, 738 s, *τ*_*uncorr*_ = 1, 721 s, *τ*_*comp*_ = 28, 551 s). **F**. Distributions of responding probability (left) and median delay (right) of followers in the same assembly as the source neuron (blue), in the next assembly (orange), or in other assemblies (gray) (*n* = 116 sequences, *n*_*within*_ = 57, *n*_*next*_ = 107, *n*_*other*_ = 326). **G**. Ratio of strong connections within assemblies, from one assembly to the next, and anywhere else, in a network first trained with unstructured inputs and then with temporally structured inputs (*n* = 10; mean±std). **H**. Same as **G** but network was first trained with one pattern and then with a different one (*n* = 10; mean±std).

Strong connections within the same assembly and from one assembly to the next were more stable than the strong connections in a network trained with unstructured Poisson inputs (Fig. 6E, blue and gray). As in the case of the assemblies in the previous section, any strong connection that turned over in a consecutive assembly pair was quickly compensated by the appearance of a new strong connection in same the assembly pair. The effective turnover rate therefore was very low, with a time constant longer than our simulations (Fig. 6E, pink). This highly stable connectivity resulted in highly responsive and stable followers: followers in the same assemblies as their source neuron or in the next assembly had a significantly higher responding probability than any follower in the other assemblies (0.64 ± 0.40 vs. 0.40 ± 0.36 for same-assembly; 0.53 ± 0.39 vs. 0.40 ± 0.36 for next-assembly), as well as shorter delays (10.5 ± 9.9 ms vs. 28.0 ± 13.4 ms for same-assembly; 13.7 ± 8.3 ms vs. 28.0 ± 13.4 ms for next-assembly), (Fig. 6F). These followers suggest that, under temporally structured inputs, our model network developed spiking sequences that could link multiple assemblies.

Finally, we tested if temporally structured inputs could flexibly reorganize networks with two types of preexisting structure: that resulting from uncorrelated inputs (Fig. 6G), and that resulting from a different pattern of temporally structured inputs (Fig. 6H). After reaching stability in the two cases, we applied a new temporally structured input pattern. In both cases, the network reorganized its strong E-to-E connections to the assembly structure described above (Fig. 6C), resulting in sequences that activated over multiple assemblies and reflecting the ordering of the new inputs.

In summary, temporally structured inputs could shape stable sequences across neural assemblies that reflect the temporal order of the inputs. The resulting strong connections had a very low turnover ratio, creating a stable backbone of propagation that sequentially linked multiple groups of neurons. Nonetheless, this stable configuration was also flexible enough to adapt to changes in the input patterns.

## Discussion

Here, we investigated how repeating sequences of spikes may arise and change under ongoing synaptic plasticity in a spiking neural network composed of recurrently connected excitatory and inhibitory neurons with biologically-plausible synaptic plasticity rules. We found the network spontaneously reached long-tailed weight distribution and structures of connectivity that made it produce sequences even in the complete absence of structured stimuli (Fig. 1). Although the statistics of these structures were stable, particular synaptic weights and sequences evolved due to the constant turnover of connections (Fig. 2, Fig. 4). Still, the timescales of turnover were longer for features of connectivity that supported sequences when compared to random connectivity (Fig. 2, Fig. 3). This lower volatility could be further reduced when the network was subjected to external structured stimuli (Fig. 5, Fig. 6). While these structured inputs reinforced connectivity structures underlying sequences, the networks retained sufficient flexibility to adapt to changes in the input (Fig. 5, Fig. 6).

Our results demonstrate that spiking sequences can be generated in randomly connected networks trained by synaptic plasticity even under unstructured inputs, which supports STDP being the main actor, while stabilizing mechanisms such as weight normalization and intrinsic plasticity [46, 59, 61] play a complementary role. Many models using STDP to produce sequences were based on binary neurons and time-discretized STDP that only considered the activity in two consecutive timesteps at a time [37, 64, 76]. This led to an idealized form of sequential activity known as “synfire chains”, and weights that binarized by saturating or decaying to zero. Other models relied on structured inputs [74] or supervised learning rules and external teaching signals [55, 56, 76, 77] which cannot account for the spontaneous sequences considered in our work and found experimentally [5, 8]. Finally, some studies are based on feedforward architectures [78, 79] or extremely small networks (fewer than 20 neurons) [74], which differ significantly from the highly recurrent large-scale networks found in the brain. The plasticity rules that we combined gave rise to stable, continuous, long-tailed weight distributions of excitatory connections that generate spiking sequences (Fig. 1, [37, 39, 57]). This particular combination of plasticity rules avoids the presence of dominant excitatory neurons that concentrate all strong connections (i.e., hub neurons found in previous work [71–73, 80]). Instead, strong connections are spread across the network, enabling sequences that could start almost everywhere (Fig. 1). These strong connections form over-represented connectivity motifs that promote sequential activation (fan-in/out motifs), consistent with experimental evidence on the routing of neuronal activity [65].

Beyond excitation, our work also involved plasticity of I-to-E connections through a symmetric iSTDP rule, resulting in a balance of excitation and inhibition (Fig. 1, [38]). In consequence, excitatory neurons that initiate sequences receive strong feedback inhibition after the onset of the sequence (Fig. S6), as observed experimentally [8]. This is the result of inhibitory neurons that start to fire after multiple excitatory neurons in the sequence have been activated. (Fig. 1). By countering these activations, this form of negative feedback mechanism might limit the growth of excitatory connections and contribute to their eventual turnover [81]. E-to-I connections were modeled as non-plastic, since their plasticity rules are rarely characterized in experiments and E-to-E and I-to-E plasticity alone was sufficient to generate sequences.

The stability of these long-tailed distributions of excitatory connections neither depended on the presence of structured input, nor it was impaired by it. Our simulations with spatially and temporally structured inputs did not affect the statistics of our excitatory weight distributions (Fig. S11). The presence of spatially structured inputs resulted in additional assembly structure within the connectivity matrix (Fig. 5, [39, 40, 42, 43, 51, 74]), within which spiking sequences persisted. Fast inputs with temporal structure, on the other hand, produced cross-assembly connections similar to “neuronal clocks” (Fig. 6, [55]). In both cases, we observed spiking sequences of reliable followers concentrated within each assembly or across two assemblies, and a slow turnover of the connectivity that produces them, highlighting the robustness and flexibility of these plasticity rules.

Long-tailed distributions of synaptic weights and sequences are widely observed phenomena across multiple brain areas and species [63,64,66,71,80–85]. Theoretical studies with different degrees of biological realism have proposed links between these weight distributions and activity patterns [37, 39, 57, 86,87]. Our results under unstructured or structured input contribute to this link and propose the role of specific plasticity rules to explain the widespread presence of both, long-tailed distributions and spiking sequences, in the brain.

Spiking sequences have been proposed to be an intrinsic feature of the brain, rather than a reflection of sequential input [4]. The brain can use these structures during learning to anchor stimulus representations. Our results demonstrating the generation of sequences even with unstructured input provide a mechanism for the emergence of this representational substrate. However, despite the robust statistics of our networks, we found a constant turnover of synaptic connections (Fig. 2) that resulted in a constant turnover of followers (Fig. 4) and thus the eventual alteration of any given sequence. If sequences are often observed in sensory, motor and navigational cortices, and are meant to bind representations, how might this turnover affect the encoding of external stimuli, actions or places?

An increasing body of experimental evidence suggests that representations in the brain might be subject to a slow “drift”, where patterns of neuronal activity correlate with task features in the short timescale, but the overall representations radically change in the long term [9]. Drift has been reported in multiple brain regions, such as mice posterior parietal cortex (PPC, [85]), mice visual cortex [11, 12], the hippocampus of mice and rats [18, 88], mice olfactory cortex [89], as well as *Drosophila* mushroom body [90]. The sequences that we describe here present sudden drops of single followers but overall slow changes of the whole sequence (Fig. 4) and thus seem suitable candidates to underlie some of these drifting representations. Recent work in the hippocampus suggests that slow drift of place representation at the scale of population codes may be the reflection of sudden changes of place fields at the level of single neurons, which is very consistent with our modeling results [91].

To deal with how a downstream area might decode a drifting representation, theoretical studies propose plastic readouts that can adjust to the encoding as long as the timescale of the drift is slower than the timescale of the decoding [92, 93]. Our sequences present a drift of the neurons involved at multiple timescales: followers late in the sequence drop out more quickly while followers earlier in the sequence are more stable (Fig. 4), resulting in a slow drift in the representation at the scale of the whole sequence. Furthermore, since most neurons in our networks could elicit a sequence, multiple sequences may run in parallel [57], which could extend even further the stability of the representations when considering the whole population. Our work with structured input strongly supports this possibility: when considering any strong connection among the followers within an assembly, turnover timescales became longer than our simulations (Fig. 5, Fig. 6).

Interestingly, in the mouse primary visual cortex, the representations corresponding to some simple and basic visual stimuli (e.g., gratings) are very stable, while the representations of more natural and complex visual stimuli are more unstable [12]. It is possible that gratings act like our temporally structured inputs, fixing neuronal representations through stereotyped repetition, while the high variance of naturalistic movies acts like our uncorrelated input, resulting in faster turnover that translates into unstable representations. A more interesting hypothesis is that such complexity-dependent drift results from the turnover dynamics we observe between early (stable) followers and late (unstable) followers (Fig. 4). Previous modeling work showed that contextual network activation determines the dynamic activation of later followers, suggesting a higher degree of selectivity of later than earlier followers [57]. Our results would thus be consistent with early neurons in the sequence representing simpler stimuli, possibly receiving direct thalamic inputs, while later neurons representing more complex stimuli. These later, complex representations would be more amenable to drift because they rely on the activation (and stability) of multiple strong connections within the recurrent circuit (Fig. 4). Since natural environments likely present a large variety of slightly different stimuli, later followers with very high stimulus selectivity may activate very rarely, and, consequently, may be more prone to turnover under our plasticity rules, resulting in a stronger drift in terms of representation. This view aligns well with prior theoretical frameworks of sparse coding, which aimed to explain why the large majority of neurons are mostly silent [94].

Although representations of sequential activity primarily rely on strong connections, weak connections may also be highly relevant for their turnover driven by plasticity. The high volatility of weak connections in our simulations (Fig. 2) suggests a quick mechanism for network computations: If weak connections are particularly susceptible to plastic changes due to the network dynamics, they may be ideally positioned to implement fast flexibility into the network. Indeed, weak connections are likely to play a role in implementing competition and cooperation across multiple sequences [57]. Under this view, reactivation of sequences under changing inputs will be first subject to the modulation of weak connectivity and may, on a longer timescale, result in the reorganization of the more stable backbone of strong connections.

Finally, although our model network may capture features of stimulus representational drift, it has a limited memory capacity. Indeed, while changing of input patterns can be seen as a feature of flexible adaptation to changing stimulus statistics (Fig. 5, Fig. 6), it can also be interpreted as a volatile memory issue, commonly known as “catastrophic forgetting” [95]. A possible avenue of future research to address this issue may lie in determining the suitable dynamics of learning of a readout layer, as described above [92, 93], or exposing the network to multiple alternating input patterns during training, similar to previous modeling work on hippocampal sequence replay [96].

## Methods

### Single neuron model

We used the adaptive exponential integrate-and-fire neuronal model (AdEx, [58]) for both excitatory and inhibitory neurons, with membrane potential dynamics:

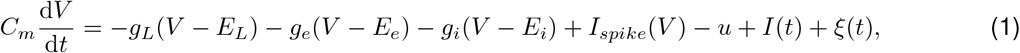

where *C*_*m*_ is the membrane capacitance, *g*_*L*_ is the leaky conductance, *g*_*e*_ and *g*_*i*_ are the conductances of the excitatory and inhibitory channels. *E*_*L*_, *E*_*e*_ and *E*_*i*_ are reversal potentials of the leaky, excitatory, and inhibitory channels, respectively (Table 1). Membrane noise is captured by *I*(*t*), a background current that follows a Gaussian distribution with mean *µ*_*wn*_ and standard deviation *σ*_*wn*_ and sampled at interval *τ*_*wn*_. Spontaneous activity and inputs from other parts of the brain not included in the model are captured by a Poisson input *ξ*(*t*), formulated as a sum of Dirac’s delta functions with amplitude *ξ*_*ext*_:

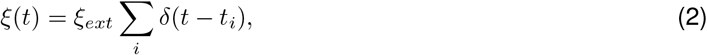

where time points {*t*_*i*_} are sampled from an exponential distribution with time constant *τ*_*ext*_. *u* is an adaptation variable following the dynamics given by:

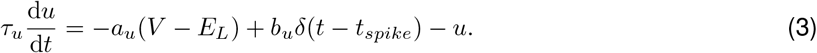

**Table 1.** Parameters for the single neuron model.

| Parameter | Description | Value |
| --- | --- | --- |
| $C_m$ | Membrane capacitance | 240 pF |
| $g_L$ | Leak conductance | 4.19 nS |
| $E_L$ | Reversal potential of the leak channel | -70.6 mV |
| $E_e$ | Reversal potential of the excitatory channel | 10 mV |
| $E_i$ | Reversal potential of the inhibitory channel | -75 mV |
| $\mu_{wn}$ | Mean of the background inputs | 62.5 pA |
| $\sigma_{wn}$ | Standard deviation of the background inputs | 21.4 pA |
| $\tau_{wn}$ | Sampling interval of the background input | 1 ms |
| $\xi_{ext}$ | Amplitude of the external Poisson input | 1 mV |
| $\tau_{ext}$ | Time constant of the external Poisson input | 3 ms |
| $a_u$ | Conductance of the adaptive channel | 4 nS |
| $b_u$ | Additional adaptive current when the neuron spikes | 80.5 pA |
| $\tau_u$ | Time constant of adaptation | 144 ms |
| $\Delta T$ | Slope constant of the spiking current $I_{spike}(V)$ | 2 ms |
| $V_{th}$ | Firing threshold | -50.4 mV |
| $V_{reset}$ | Reset potential | -60 mV |
| $V_{peak}$ | Peak potential to identify spikes | 0 mV |
| $\tau_e$ | Time constant of the excitatory channel | 1.1 ms |
| $\tau_i$ | Time constant of the inhibitory channel | 1.1 ms |
| $dt$ | Simulation time step | 0.1 ms |

*I*_*spike*_(*V* ) is an additional exponential function of the membrane potential *V* to simulate the spiking process. It can be written as

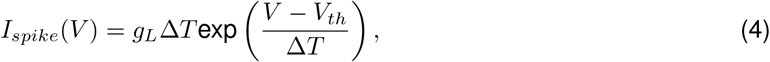

where *V*_*th*_ is the firing threshold and Δ*T* is a slope parameter which controls the steepness of the membrane potential trace just before a spike. The spike times of one neuron are determined by the membrane potential exceeding *V*_*peak*_. After a spike, the membrane potential is reset to *V*_*reset*_.

Excitatory and inhibitory conductances, *g*_*e*_ and *g*_*i*_, are determined by the weights of input synapses:

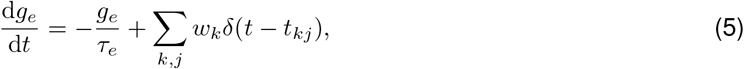

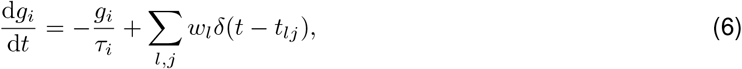

where the indices *k* and *l* correspond to presynaptic excitatory and inhibitory neurons, *w* denotes the synaptic weight, and *j* denotes the index of a single spike so that *t*_*kj*_ represents the timing of the spikes of the presynaptic neuron *k*. All parameters for the single neuron model are provided in Table 1.

### Network model

The model network consisted of *N*_*e*_ = 1200 excitatory (E) and *N*_*i*_ = 240 inhibitory (I) neurons with sparse connections between all neuron types (Fig. 1A, Table 2). Initial connectivity was random and homogeneous, with weak strengths unable to cause a postsynaptic spike with a single presynaptic one. E-to-E and I-to-E connections were plastic, while E-to-I and I-to-I connections were static. No new connections were created even if some of them decreased to zero due to plasticity. A synaptic delay of 1.5 ms was modeled for all connections.

**Table 2.** Parameters in the network model.

| Parameter | Description | Value |
| --- | --- | --- |
| $p_{ee}$ | Connection probability from E to E | 0.06 |
| $p_{ei}$ | Connection probability from E to I | 0.19 |
| $p_{ie}$ | Connection probability from I to E | 0.2 |
| $p_{ii}$ | Connection probability from I to I | 0.11 |
| $T^{ee}$ | Sum of incoming E-to-E connections | 297 nS |
| $T^{ei}$ | Sum of incoming E-to-I connections | 1012 nS |
| $T^{ie}$ | Sum of incoming I-to-E connections | 643 nS |
| $T^{ii}$ | Sum of incoming I-to-I connections | 373 nS |
| $d_{syn}$ | Synaptic delay | 1.5 ms |

### Synaptic plasticity rules

We implemented four different plasticity rules: excitatory spike-timing-dependent plasticity (STDP) for E-to-E connections, inhibitory STDP for I-to-E connections, weight normalization for E-to-E and I-to-E connections, and intrinsic plasticity for E neurons (Fig. 1A).

Spike-timing-dependent rules and intrinsic plasticity were applied immediately after a pair of spikes was detected. Weight normalization was applied whenever the sum of incoming or outgoing synaptic weights changed by more than a given threshold (1% in this study).

### Excitatory STDP (eSTDP)

The change in synaptic weight of an E-to-E connection was implemented as [46]

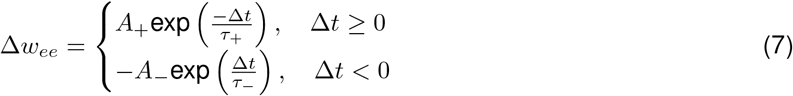

for any pair of pre- and postsynaptic excitatory spikes separated by a temporal difference of Δ*t* = *t*_*post*_ − *t*_*pre*_. *A*_+_ and *A*_−_ determine the maximum amplitude of the long-term potentiation (LTP) or long-term depression (LTD) of the synapse when Δ*t* is (close to) 0. All parameters for the plasticity rules are provided in Table 3.

**Table 3.**
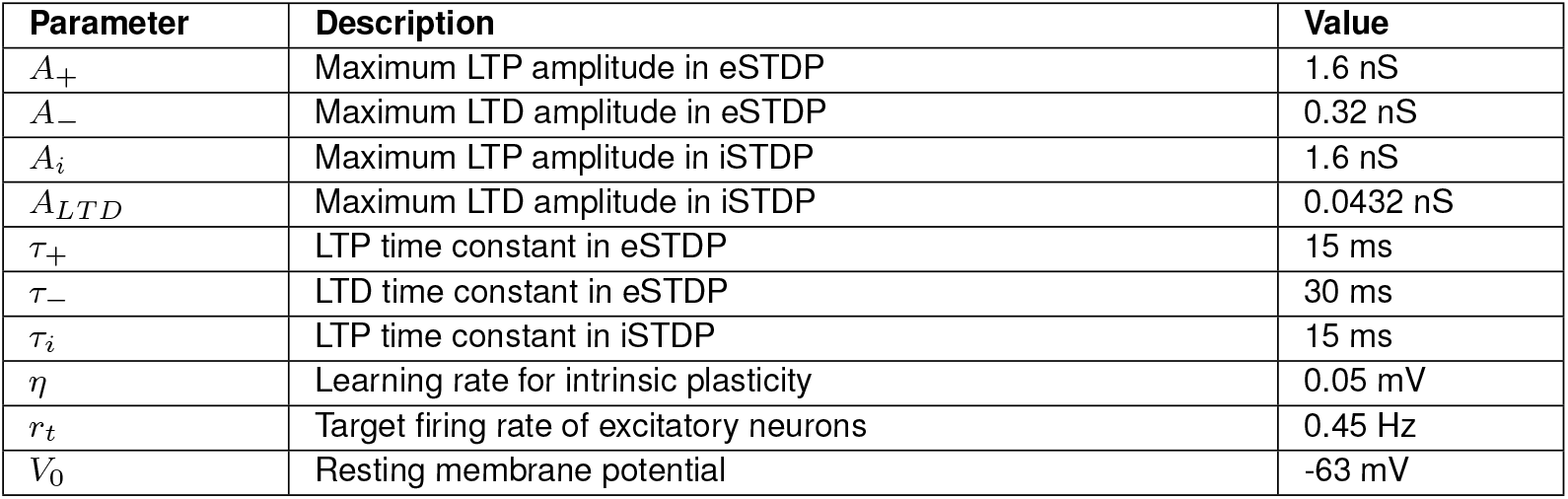
Parameters for the plasticity rules.

### Inhibitory STDP (iSTDP)

The change in synaptic weight of an I-to-E connection was determined by [38]

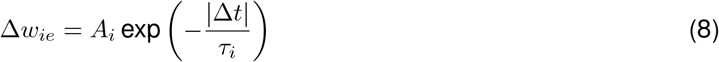

for any pair of inhibitory presynaptic and excitatory postsynaptic spikes separated by a temporal difference of Δ*t* = *t*_*post*_ − *t*_*pre*_. Additionally, whenever there was a single inhibitory presynaptic spike, independently of the presence or absence of a postsynaptic excitatory one, a corrective LTD weight change was applied:

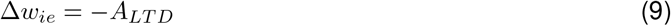

### Weight normalization

Pre- and postsynaptic weight normalization was applied to both E-to-E and I-to-E connections. Postsynaptic normalization for E-to-E connections ensured that the sum of incoming excitatory synaptic weights for each postsynaptic excitatory neuron remained constant at *T* ^*ee*^ using:

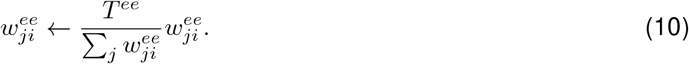

Presynaptic normalization kept the sum of outgoing synaptic weights from excitatory neuron *i* at 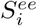, using the updating rule:

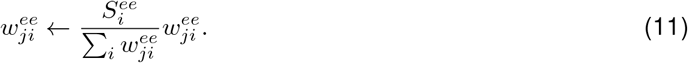

Similarly, for I-to-E connections, the sum of incoming inhibitory synaptic weights for each postsynaptic excitatory neuron was kept at *T* ^*ie*^, while the sum of outgoing synaptic weights from inhibitory neuron *i* remained constant at 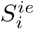, using the rules:

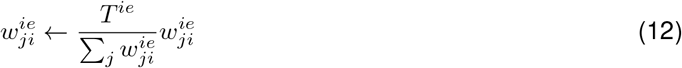

and

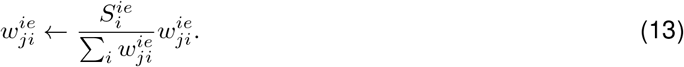

### Intrinsic plasticity for excitatory neurons

The firing threshold *V*_*th*_ of each E neuron was adjusted to match a universal target firing rate *r*_*t*_:

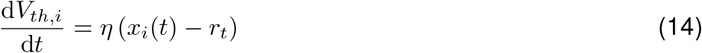

where *V*_*th,i*_ is the firing threshold of neuron *i, x*_*i*_(*t*) is equal to 1 if the neuron spikes at time *t* and otherwise to 0 (with units of Hz). The parameter *η* controls the speed of this adjustment.

### Effective weights

Due to the dynamic firing threshold of excitatory neurons and in order to quantify the real impact of synapses, we defined the “effective weight” as:

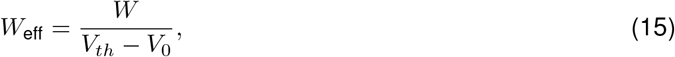

that is, the ratio of the synaptic weight *W* to the voltage distance from the resting potential *V*_0_ to the firing threshold *V*_*th*_. The resting potential *V*_0_ was estimated through simulation of the membrane potential with only the background input *I*(*t*).

### Alternative combinations of plasticity rules

To evaluate the contribution of each plasticity rule, we conducted simulations where one or two of these rules was disabled (Fig. S12). Removing both presynaptic and postsynaptic normalization led to runaway dynamics of the network, while all other combinations resulted in steady states with approximately lognormal E-to-E weight distributions (Fig. S12A-B).

We approximated the proportion of neurons capable of initiating a sequence as the proportion of excitatory neurons with maximal outgoing *W*_eff_ greater than 2.5 nS/mV (Fig. S2A). In the full plasticity model, this approximated proportion was 90.5% (Fig. S12C, blue), closely matching the actual proportion of 87.1% of sequence-generating neurons observed through direct testing (n=155 neurons, Fig. S2). Using this approximation, we could assess the effectiveness of the stable rule combinations:

1. Removing only intrinsic plasticity: 2.25%(Fig. S12C, orange).
2. Removing only presynaptic normalization: 21.42% (Fig. S12C pink).
3. Retaining only postsynaptic normalization: 3.08% (hub neurons, Fig. S8, Fig. S12C, cyan).
4. Retaining only presynaptic normalization: 70.17% (Fig. S12C, purple).
5. Retaining only presynaptic normalization and intrinsic plasticity: 96.08%.

Interestingly, combinations 4 and 5, which rely on presynaptic normalization, came close and even surpassed the full model. However, using only presynaptic normalization caused the network’s firing rate to increase over time, ultimately far exceeding biologically observed rates (Fig. S12D) [8]. While intrinsic plasticity can regulate the firing rate in such scenarios, it drives the firing threshold up to -18 mV, which is not biologically realistic. For these reasons, we used the full plasticity model in our simulations, even though presynaptic normalization alone might be sufficient to enable sequence generation.

### Sequence testing

We tested for the presence of sequences similarly to previous experimental and modeling studies [8, 57]. For the testing procedure below, we made a frozen copy of the network with disabled plasticity.

We artificially triggered a series of spikes in a randomly selected neuron, which we defined as the “source neuron”of the sequence. Then we calculated the “firing rate modulation” of all other neurons in the network, defined as the change in their firing rates before and after the source neuron’s spike. Neurons exhibiting a firing rate modulation significantly higher than expected under a statistical null model were identified as “followers” in the sequence.

The same source neuron was forced to spike every 400 ms over *M* = 1, 000 trials (Fig. 1E). To compensate for this additional forced activity, the frequency of the external Poisson input *ξ*(*t*) was reduced (*τ*_*ext*_ = 4 ms). In each trial and for each neuron in the network, we computed *r*_1_ as the firing rate in the 100 ms before the forced spike (Fig. 1E, gray shades) and *r*_2_ as the firing rate in the 300 ms after (Fig. 1E, blue shades).

To construct a statistical null model, we calculated the averages of *r*_1_ and *r*_2_ (⟨*r*_1_⟩ and ⟨*r*_2_⟩) across all neurons and trials, and used them to simulate a single neuron firing Poisson spikes (instead of following Eq. 1) over *M* ^*′*^ = 10, 000 trials. Hence, in each trial, this idealized neuron was set to fire at rate ⟨*r*_1_⟩ for the first 100 ms and at rate ⟨*r*_2_⟩ in the next 300 ms.

We defined the “firing rate modulation” of each neuron Δ*r* as Δ*r* = *r*_2_ − *r*_1_. Each pair of *r*_1_ and *r*_2_, for each trial, contributed one observation of Δ*r*. To reduce sampling noise, we divided the *M* = 1, 000 simulated observations and the *M* ^*′*^ = 10, 000 null observations of Δ*r* into 10 equal-sized subsets (100 simulated and 1,000 null observations for each). In each subset, we computed the 95th percentile of Δ*r* from the null model. A neuron was identified as a follower only if its average ⟨Δ*r*⟩ exceeded the 95th percentile of the null model in all of the 10 subsets.

To calculate the responding probability, delay, and jitter(Fig. S4C-D), for every identified follower, in each of the *M* = 1, 000 trials of spike triggering, we searched for its first spike after the forced source spike (Fig. 1E). If none was detected within a 300 ms window after the spike triggering, we considered the follower unresponsive to the source neuron for that trial. If a spike was found, the elapsed time from the triggered spike was recorded as the delay. If a follower responded to the source neuron in *M* ^∗^ out of the *M* = 1, 000 trials, its responding probability was then *M* ^∗^*/M* (Fig. 1E). A total of *M* ^∗^ delays were recorded, with the median calculated as the median delay, and the standard deviation defined as the jitter.

After extracting the sequences generated from all excitatory neurons, we estimated the minimal effective weight required to trigger a follower. The threshold was estimated to be 2.5 nS/mV (Fig. S2C) and determined as the criterion to classify strong connections.

### Turnover of connections

We calculated the turnover of existing connections by estimating the mean proportion that remains at a given time interval (Fig. 2D). We used up to 11 evenly-spaced snapshots of the network after it had reached steady state. For example, to estimate turnover of connections at a 2, 000*s* interval for a simulation that reached steady state at 8, 000*s* and lasted a total of 16, 000*s*, we took snapshots at *t* = 8, 000*s*, 10, 000*s*, 12, 000*s*, 14, 000*s*, 16, 000*s*. Then, we calculated the ratio of connections that remained from one snapshot to the next and averaged them across all snapshots, resulting in one estimate per instantiation of the network.

Pooling estimates across multiple instantiations and multiple time intervals, we estimated the turnover time constant *τ* of connections by fitting using an exponential function:

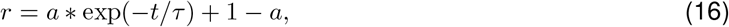

where *r* is the remaining ratio and *t* is the time interval. If there was no baseline in the exponential function (Fig. 2D, 5E, 6E), we fixed *a* = 1 in the fitting.

The turnover of motifs were calculated following the exact same procedure.

### Motif analysis

The frequency of motifs composed of strong E-to-E connections (*F*, Fig. 3C colored) was defined as the ratio of the total count of motifs (*N* ) and the number of all potential excitatory motifs that could exist in the network (*N*_0_):

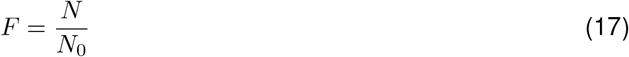

*N*_0_ is different for each motif type and can be calculated from the connection probability (*p*_*ee*_) and the number of excitatory neurons (*N*_*e*_).

For linear chains,

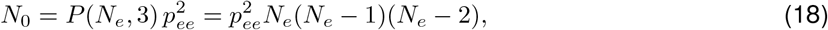

where *P* is the number of permutations. For convergence and divergence motifs,

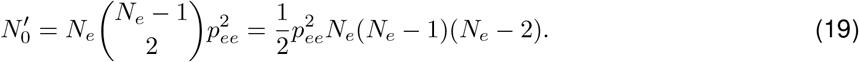

For fan-in/out motif,

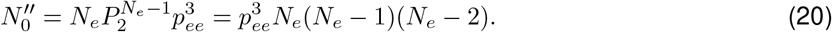

We compared this frequency (*F* ) with the one expected if strong connections were randomly shuffled (*F*_*rand*_, Fig. 3C black). We use the ratio (*p*_0_, Fig. 2B) of strong E-to-E connections to the expected number of E-to-E connections (*p*_*ee*_*N*_*e*_(*N*_*e*_ − 1)) to calculate the expected count of a particular motif in a fully random network (*N*_*rand*_). For linear chains,

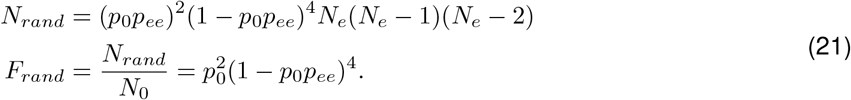

For convergence and divergence motifs,

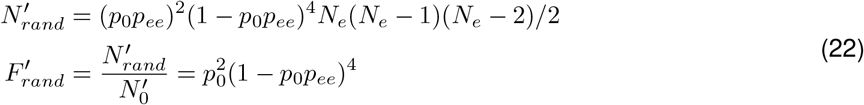

For fan-in/out motif,

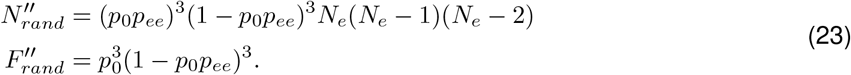

### Follower turnover

When analyzing follower turnover based on responding probabilities, delays, and synaptic jumps (Fig. 4D, 4E, and S10A), we selected an interval of 1,000 s and took 9 snapshots of the network from *t* = 8, 000 s to *t* = 16, 000 s, generating 8 intervals to calculate the remaining ratio of followers. Taking Fig. 4D as an example, we selected 50 sequences and went over these 8 intervals, calculating the remaining ratio of the followers with given responding probability. This yielded 400 ratios, which were averaged using the number of followers in each snapshot as weights.

### Spatially and temporally structured inputs

To simulate spatially structured inputs, we introduced an input pattern with pairwise correlations to the network, superimposed on the unstructured Poisson input as a background. The pattern was composed of 12 randomly partitioned assemblies of 100 excitatory neurons each. These assemblies were sequentially activated with correlated inputs for 100 s, separated by 5 s gaps between activations (Fig. 5A, also see table below for parameters). At any given time, only one assembly received correlated inputs, while all other assemblies and inhibitory neurons were driven by unstructured Poisson inputs. Note that, when switching from one pattern to another (Fig. 5H), since patterns were random and all neurons belonged to some assembly, the two patterns had a slight overlap. The network reached a steady state after ∼10,000 s of training (Fig. 5B-C). All parameters for the spatially structured inputs are provided in Table 4.

**Table 4.** Parameters of the spatially structured inputs (Fig. 5)

| Parameter | Description | Value |
| --- | --- | --- |
| $T_{corr}$ | Period in which correlated inputs are given to an assembly | 100 s |
| $T_{gap}$ | Period in which no correlated inputs are given (between two periods with correlation) | 5 s |
| $\rho_{corr}$ | Correlation of correlated external inputs | 1.0 (fully correlated) |
| $\xi_{ext,e}$ | Amplitude of external inputs to E neurons | 20 mV |
| $\tau_{ext,e}$ | Time constant of external inputs to E neurons | 400 ms |
| $\xi_{ext,i}$ | Amplitude of external inputs to I neurons | 20 mV |
| $\tau_{ext,i}$ | Time constant of external inputs to I neurons | 300 ms |

For generating temporally structured inputs, we modeled inputs as Poisson spikes targeting one assembly at a time, then shifted sequentially to the next. Each assembly was activated for 10 ms and then received no input until the next cycle. Inhibitory neurons still received unstructured Poisson inputs at any time (also see table below for parameters). All parameters for the temporally structured inputs are provided in Table 5.

**Table 5.**
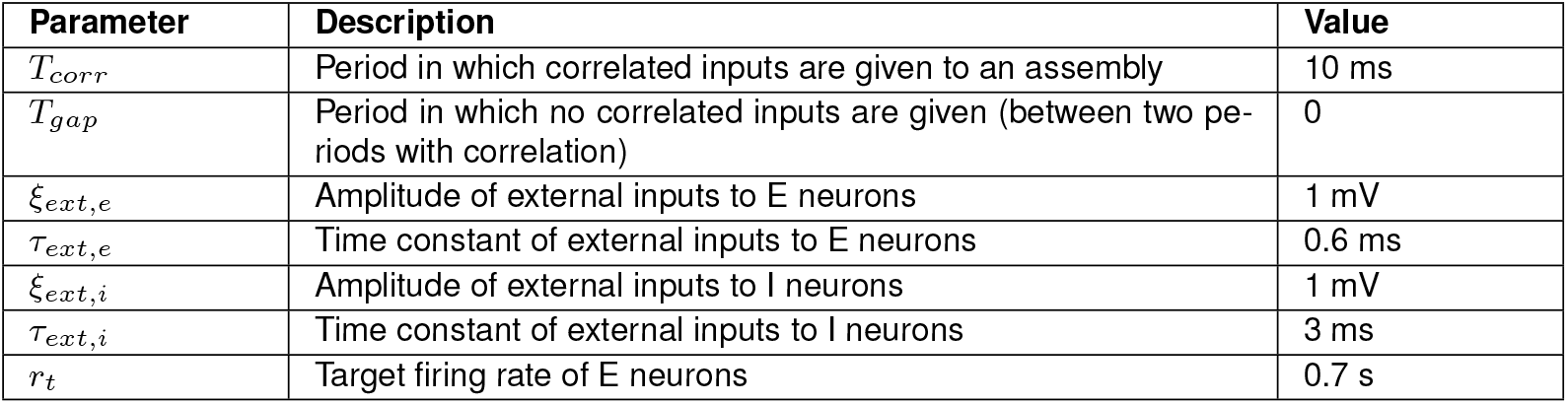
Parameters of the temporally structured inputs (Fig. 6)

| Parameter | Description | Value |
| --- | --- | --- |
| $T_{corr}$ | Period in which correlated inputs are given to an assembly | 10 ms |
| $T_{gap}$ | Period in which no correlated inputs are given (between two periods with correlation) | 0 |
| $\xi_{ext,e}$ | Amplitude of external inputs to E neurons | 1 mV |
| $\tau_{ext,e}$ | Time constant of external inputs to E neurons | 0.6 ms |
| $\xi_{ext,i}$ | Amplitude of external inputs to I neurons | 1 mV |
| $\tau_{ext,i}$ | Time constant of external inputs to I neurons | 3 ms |
| $r_t$ | Target firing rate of E neurons | 0.7 s |

### Entropy and rank correlation of the sequence

Following the same calculation as in [8], the entropy of a sequence was defined as

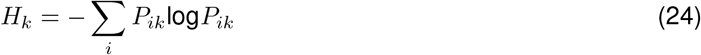

where *P*_*ik*_ is the probability for follower *i* to be the *k*^*th*^ to fire in the sequence. In a totally random sequence with *n* followers, *P*_*ik*_ = 1*/n* for all the followers and the entropy for any *k* equals to

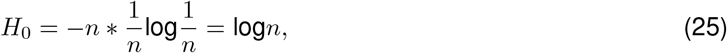

allowing us to define the normalized entropy as *H*_*k*_*/H*_0_. We took the *n* = 135 sequences in Fig. S2, calculated their normalized entropy, and then created 10 shuffled sequences for each of them (1350 in total) as a null model (Fig. S4A).

To calculate the rank correlation of sequences (Fig. S4B), we first defined an order matrix for every trial in which a spike was triggered. Specifically, for trial *k*, the order matrix *O*_*k*_(*i, j*) was defined as

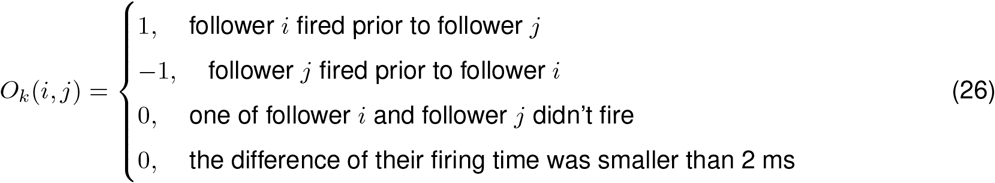

Then we defined

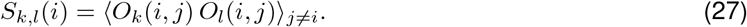

As a result, a positive *S*_*k,l*_(*i*) means the firing order of follower *i* relative to other followers was similar in trial *k* and trial *l*. A negative *S*_*k,l*_(*i*) means the firing order was reversed. Then we could calculate the rank correlation for neuron *i* as

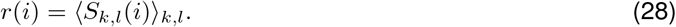

### Hub neurons

Removing presynaptic normalization and intrinsic plasticity led to the appearance of hub neurons (Fig. S8). For these simulations, some parameters were different as listed in Table 6 to avoid runaway activity through explosive excitation (STDP parameters) or to show the EPSPs triggered solely by the forced spike (*ξ*_*ext*_).

**Table 6.** Hub neurons.

| Parameter | Description | Value |
| --- | --- | --- |
| $A_+$ | LTP amplitude in eSTDP | 0.4 nS |
| $A_-$ | LTD amplitude in eSTDP | 0.08 nS |
| $A_i$ | LTP amplitude in iSTDP | 0.4 nS |
| $A_{LTD}$ | LTD amplitude in iSTDP | 0 |
| $\xi_{ext}$ | Amplitude of external Poisson inputs (only for Fig. S8D) | 0 |

## Acknowledgments

This work was supported by funding from Max Planck Society and the European Research Council (ERC) under the European Union’s Horizon 2020 research and innovation program (Grant agreement No. 804824). We thank group members of the ‘Computation in Neural Circuits’ group for discussions and readings of the manuscript.

## Code availability

All code is publicly available at:

https://github.com/comp-neural-circuits/spontaneous-emergence-neural-sequences

## Author contributions

S.S., J.L.R., and J.G. conceived the research. S.S. developed the model and performed the simulations. S.S., J.L.R., and J.G. wrote the manuscript.

## Supplementary Figures

**Figure S1.**
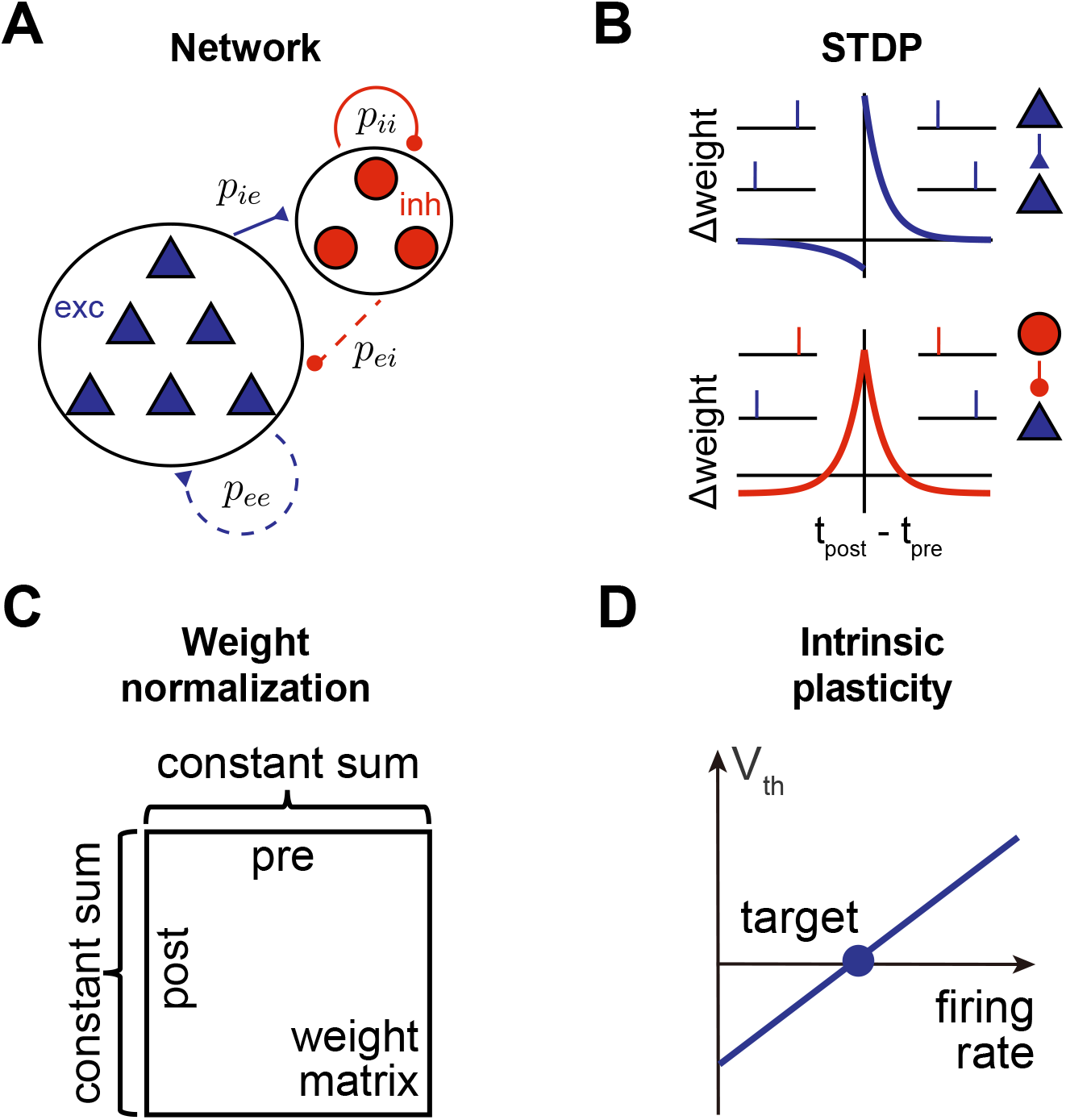
Synaptic plasticity mechanisms in the model neural network. follow a Hebbian pair-based eSTDP rule and I-to-E connections follow a pair-based iSTDP rule. Synaptic normalization preserves the total sum of incoming and outgoing weights (E-to-E and I-to-E). Intrinsic adjusts the firing threshold of excitatory neurons if the firing rate is higher or lower than the target firing rate. **A**. Schematic of the model network with 1,200 excitatory and 240 inhibitory AdEx neurons, where E-to-E and I-to-E connections are plastic. **B**. The STDP rules that changes the E-to-E (eSTDP) and I-to-E (iSTDP) synaptic weights. **C**. Synaptic normalization preserves the total sum of incoming and outgoing weights (E-to-E and I-to-E). **D**. Intrinsic plasticity adjusts the firing threshold of excitatory neurons if the firing rate is higher or lower than the target firing rate.

**Figure S2.**
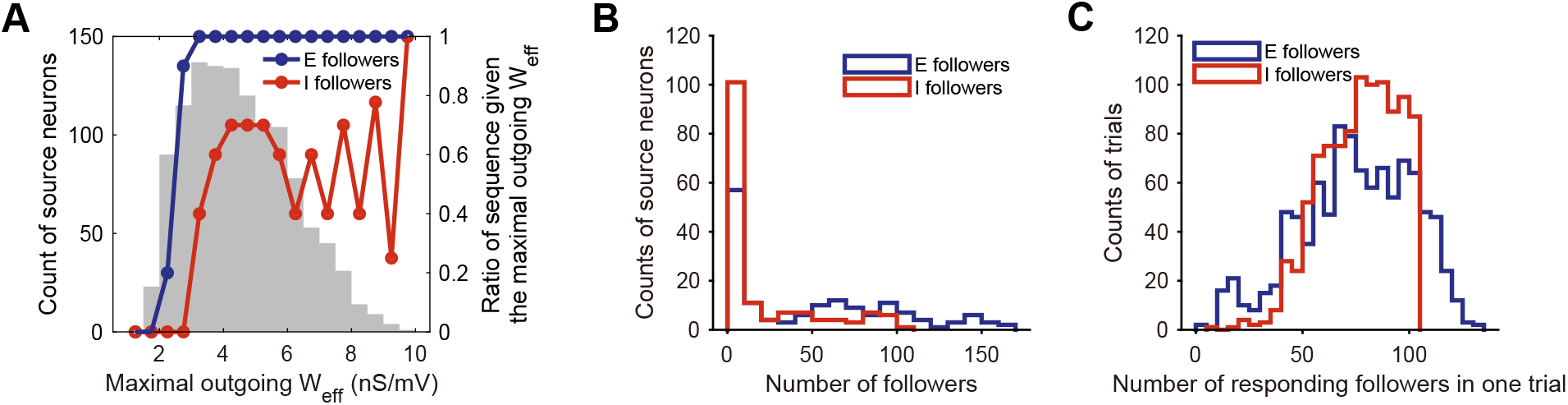
Statistics of sequence generation. **A**. Sequence generation plotted against source neurons’ maximal outgoing effective weight. (Gray bars) Histogram of the maximal outgoing *W*_eff_ of the 1,200 excitatory neurons in a model network. (Blue line) The ratio of source neurons that had at least one excitatory follower, given the maximal outgoing *W*_eff_, calculated within the *n* = 155 source neurons that we chose. (Red line) The ratio of source neurons that had at least one inhibitory follower, given the maximal outgoing *W*_eff_, calculated within the *n* = 155 source neurons that we chose. **B**. Distribution of the number of excitatory followers and inhibitory followers of the *n* = 155 source neurons that we tested. **C**. Distribution of the number of followers that spiked after the forced spike of the same source neuron. Pooled from 1,000 consecutive trials.

**Figure S3.**
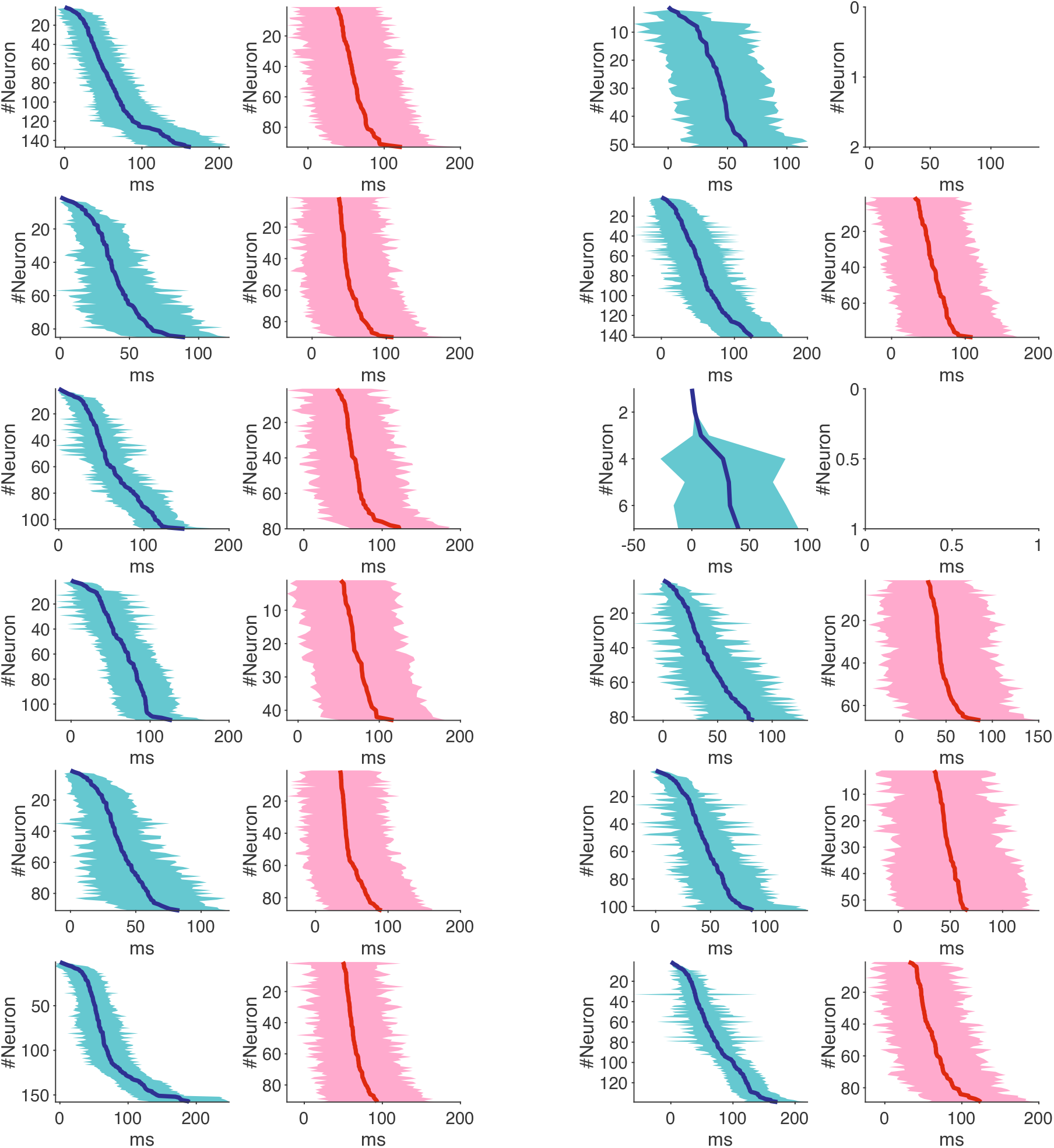
More examples of sequences generated in the same network as in Fig. 1. Blue curves indicate median delays of excitatory followers. Red curves indicate median delays of inhibitory followers. Cyan and pink shades indicate jitters.

**Figure S4.**
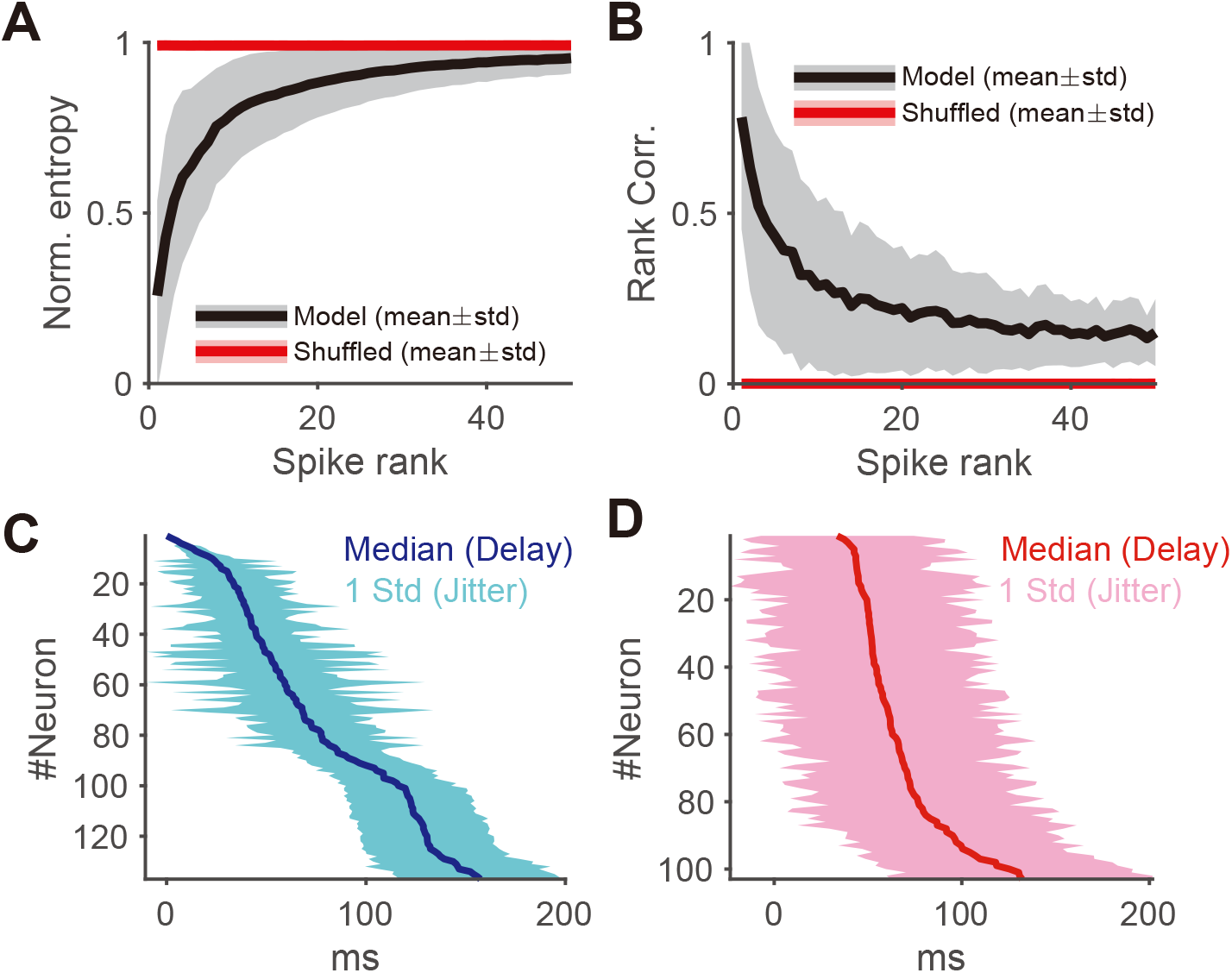
Sequence properties. **A**. Normalized entropy of sequence in the model network (black) and the null model with shuffled sequences (red). Data pooled from all source neurons with at least one follower (*n* = 135). Each sequence was shuffled 10 times. **B**. Rank correlation as a function of spike rank in the model network (black) and the null model with shuffled sequences (red). Same data as D. **C**. Delay of excitatory followers after source spike. Solid line indicates the median delay and shading indicates the jitter (standard deviation across all trials). **D**. Same as C but for inhibitory followers.

**Figure S5.**
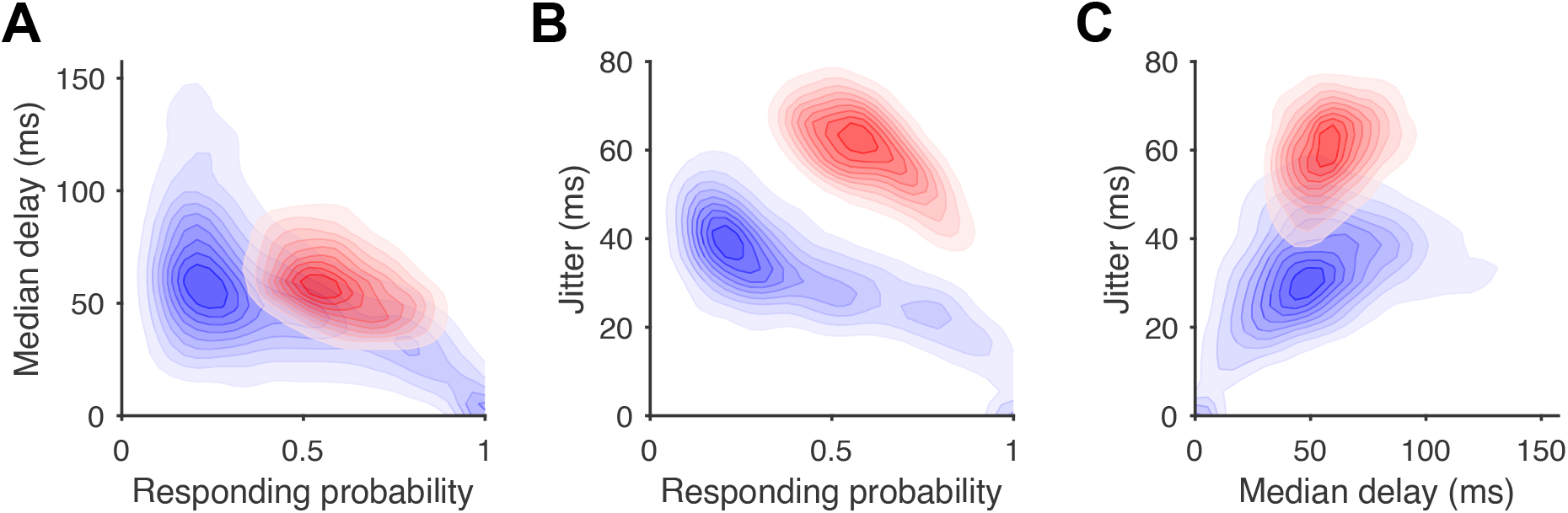
Two dimensional density plot of followers’ responding probability, median delay, and jitter. **A**. Density plot of responding probability and median delay of followers. Median delay is negatively correlated to responding probability. Pearson correlation *r* = −0.507. **B**. Density plot of responding probability and jitter of followers. Jitter is negatively correlated to responding probability. Pearson correlation *r* = −0.246. **C**. Density plot of median delay and jitter of followers. Jitter is positively correlated to median delay. (Blue) Excitatory followers; (Red) Inhibitory followers. Darker color indicates higher probability density. Pearson correlation *r* = 0.343. Pooled over 7,769 excitatory followers and 2,796 inhibitory followers from 135 sequences.

**Figure S6.**
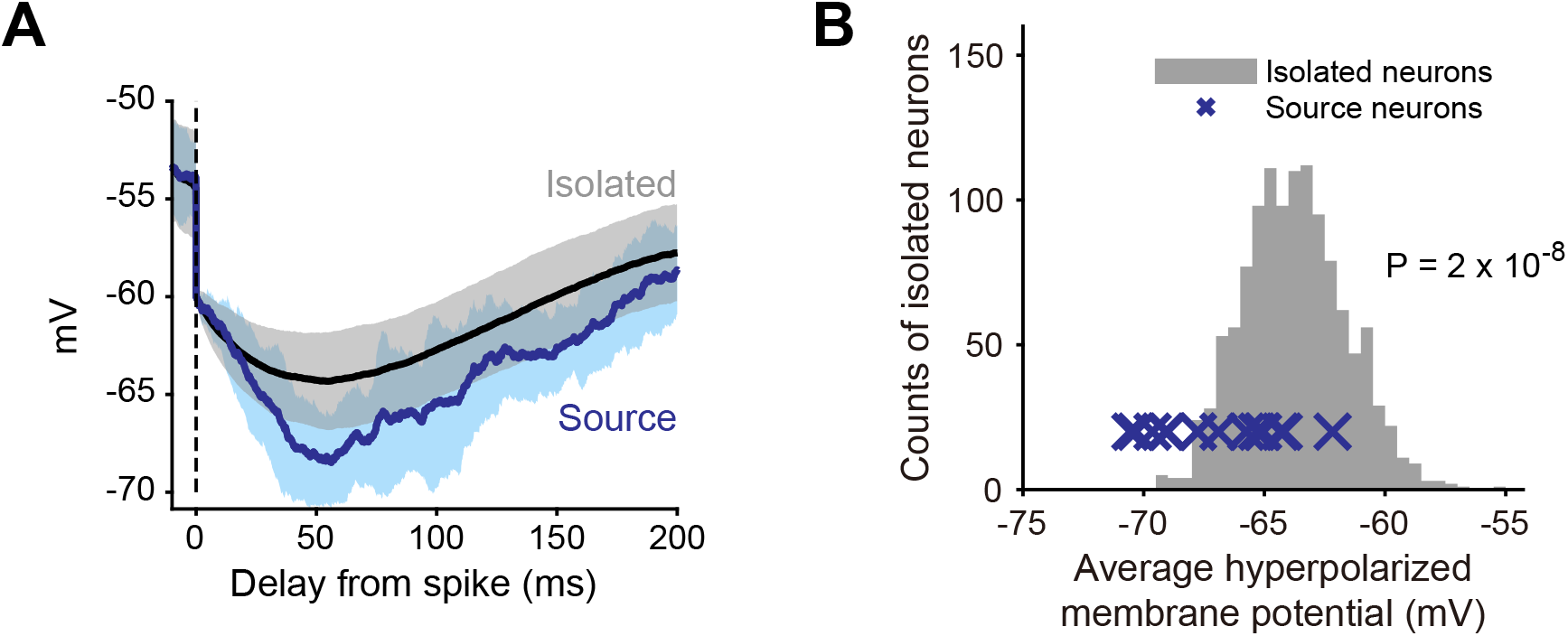
Feedback inhibition from inhibitory followers. **A**. Source neuron triggers feedback inhibition in the model. Blue: source neuron membrane potential after a spike. Pooled over *n* = 15 neurons. Black: membrane potential of isolated neurons. Pooled over *n* = 1, 200 neurons. Solid lines indicate means and shadings the standard deviations. **B**. Quantification of feedback inhibition (A). The average hyperpolarized membrane potential is quantified as the mean of the membrane potential within 50 ms to 100 ms after the spike.

**Figure S7.**
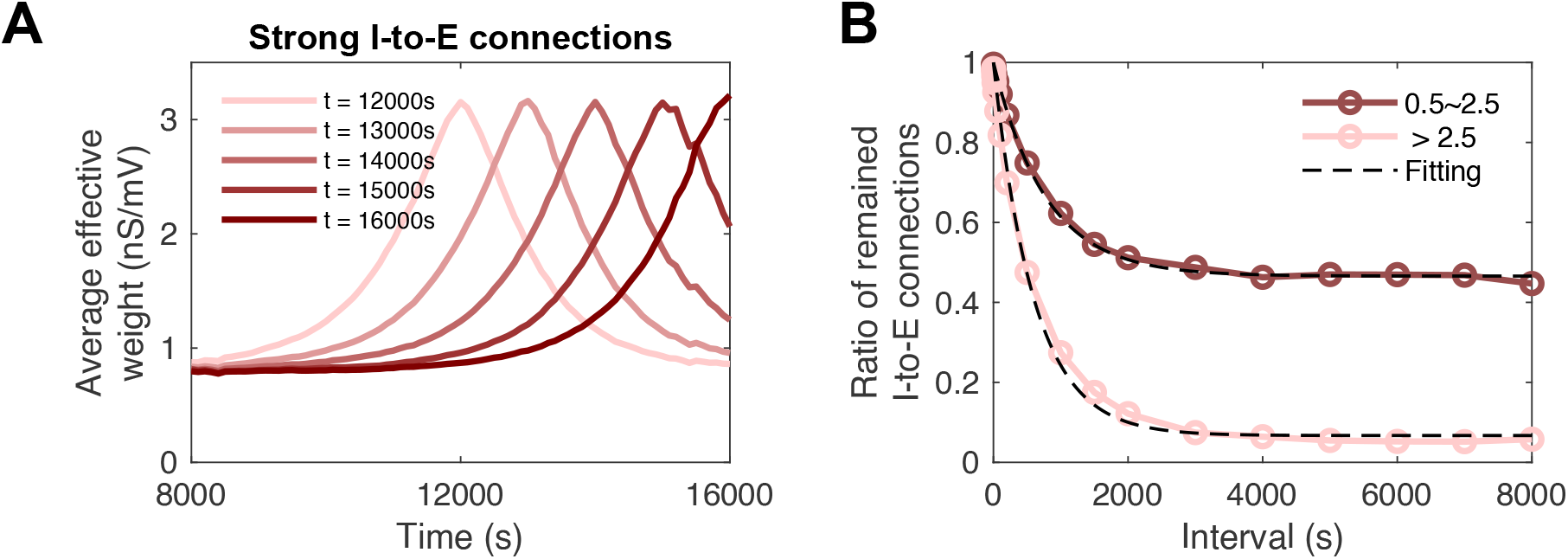
The turnover of I-to-E connections. **A**. Average strength of multiple groups of I-to-E connections picked from the tail of the *W*_eff_ distribution at different time points, after the weight distribution has reached a steady state. **B**. The decay rate of strong I-to-E weights (*W*_eff_ *>* 2.5) and weak I-to-E connections (0.5 *< W*_eff_ *<* 2.5). Dots indicate ratios calculated from simulations with 10 instantiations of the network. Dashed lines indicate exponential fits with different time constants and baselines for each group.

**Figure S8.**
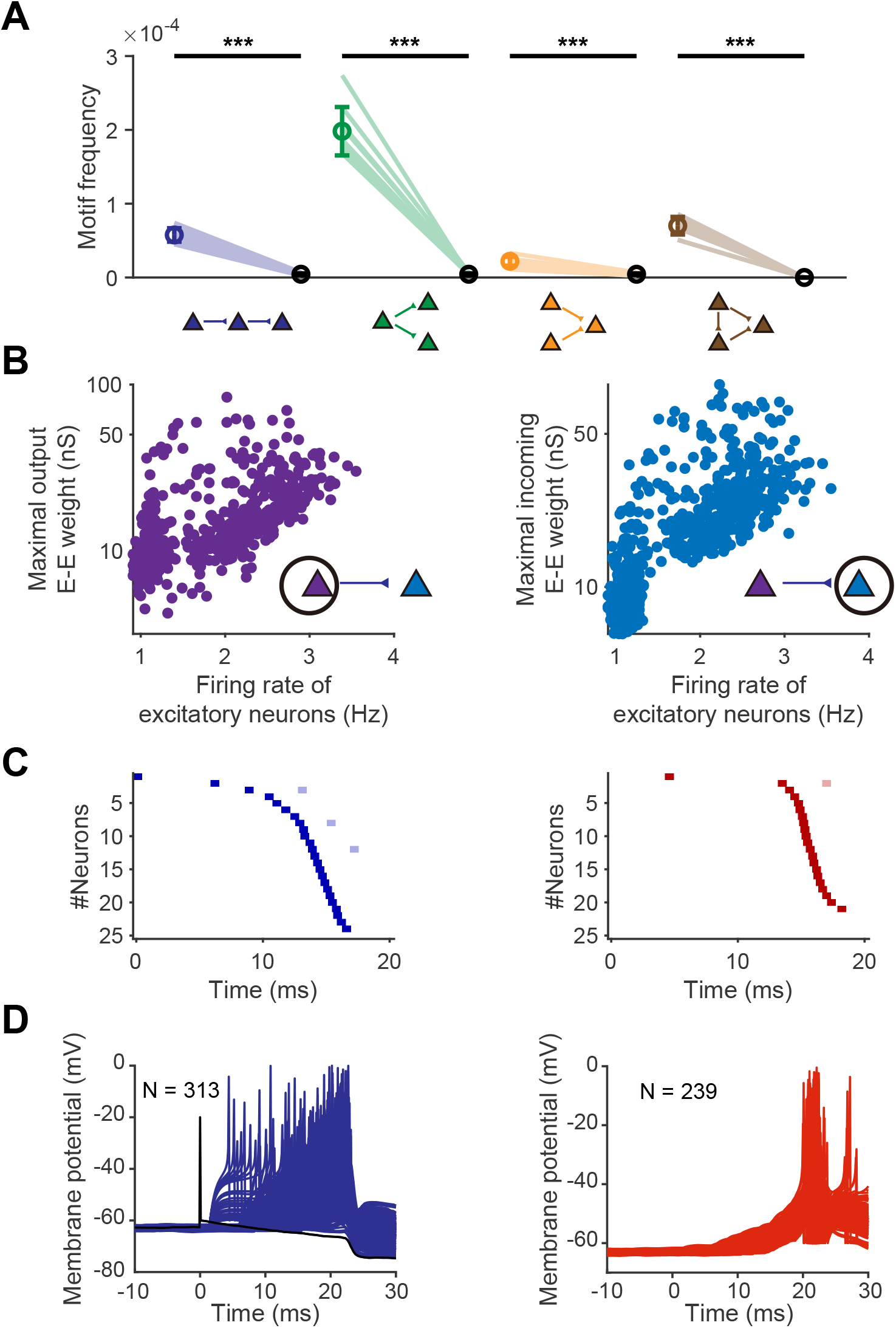
A model network without presynaptic normalization and intrinsic plasticity generates hub neurons and short sequences. **A**. The comparison between the motif frequency at *t* = 6, 000 s in the alternative model network with hub neurons and the estimated ratio in a random control network given the *p*_0_ value at that time point. Calculated from 10 trials. Error bars represent standard deviation. ***: *P <* 0.001. **B**. Hub neurons. The synaptic strength is proportional to the firing rate of neurons. Neurons with a high firing rate have a much greater impact on the network dynamics. Left: presynaptic neurons in E-to-E connections. Right: Postsynaptic neurons in E-to-E connections. **C**. Short sequences as a result of hub neurons. Left: excitatory followers. Right: inhibitory followers. **D**. Explosive dynamics in which a great number of followers can be generated within a time window as short as 30 ms. Left: excitatory neurons. Right: inhibitory neurons.

**Figure S9.**
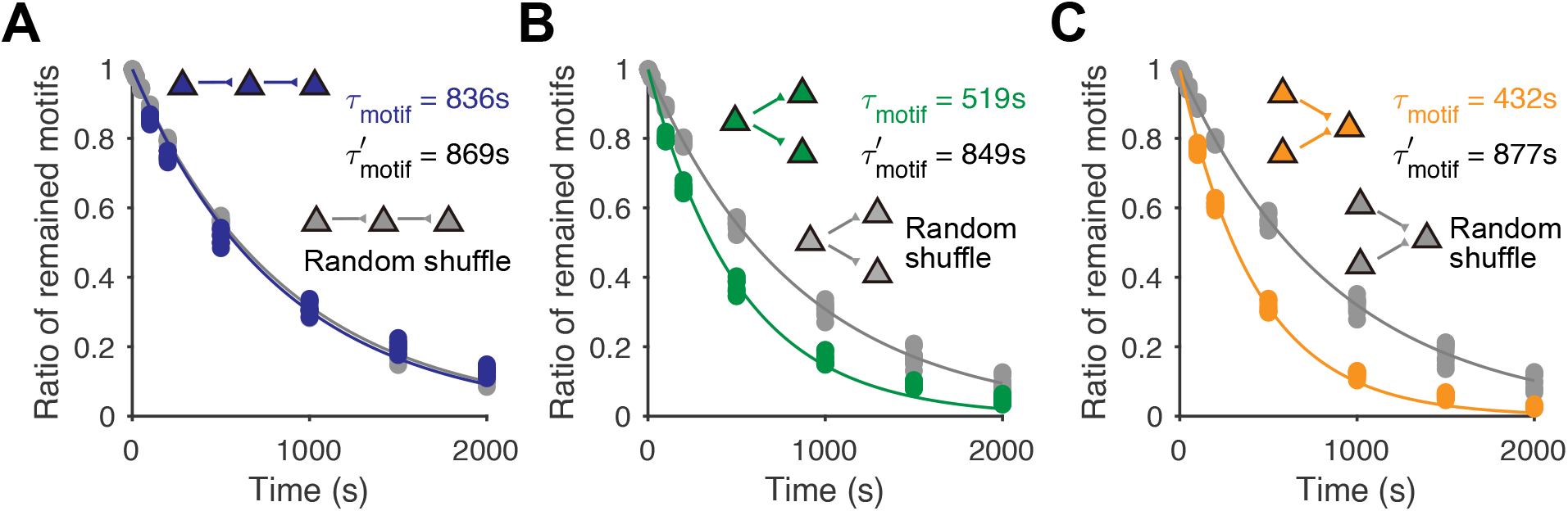
The decay of three-neuron motifs in our model network (color-coded) and in the null model (gray) in which the connections were shuffled. **A**. Linear chains. The decay of linear chains in our model network was comparable to the null model, consistent with Fig. 3D in which the ratio of linear chains was comparable to the null model. **B**. divergence motifs. The decay of divergence motifs in our model network was much faster than the null model, consistent with Fig. 3D in which the ratio of divergence motif was much lower than the null model. **C**. Convergence motifs. The decay of convergence motifs in our model network was much faster than the null model, consistent with Fig. 3D in which the ratio of convergence motifs was much lower than the null model.

**Figure S10.**
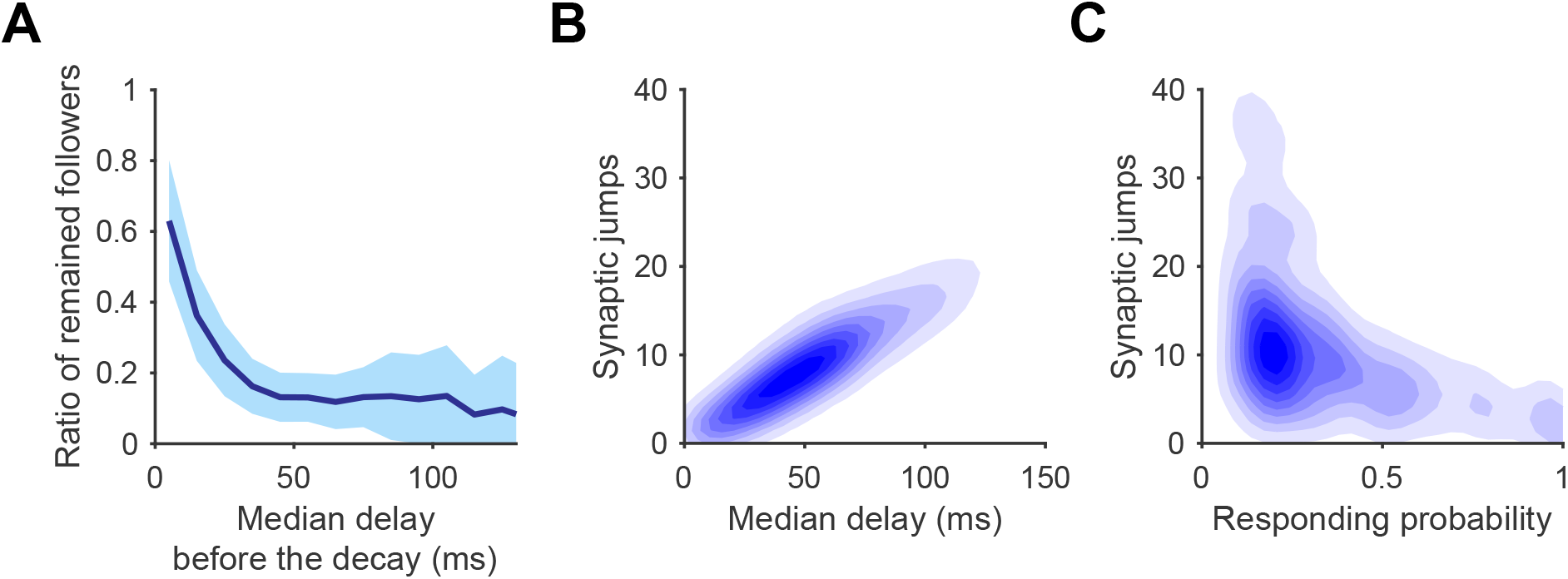
**A**. The remaining ratio of followers after a 1,000 s interval decreases with the median delay of followers. Solid line indicates the average and shade represents the standard deviation. **B**. The strong synaptic jump is negatively correlated to the responding probability. Pearson’s correlation *r* = − 0.386, pooled over 2,508 followers at *t* = 8, 000 s in Fig. 4. **C**. The strong synaptic jump is positively correlated to the median delay. Pearson’s correlation *r* = 0.494, same followers as B.

**Figure S11.**
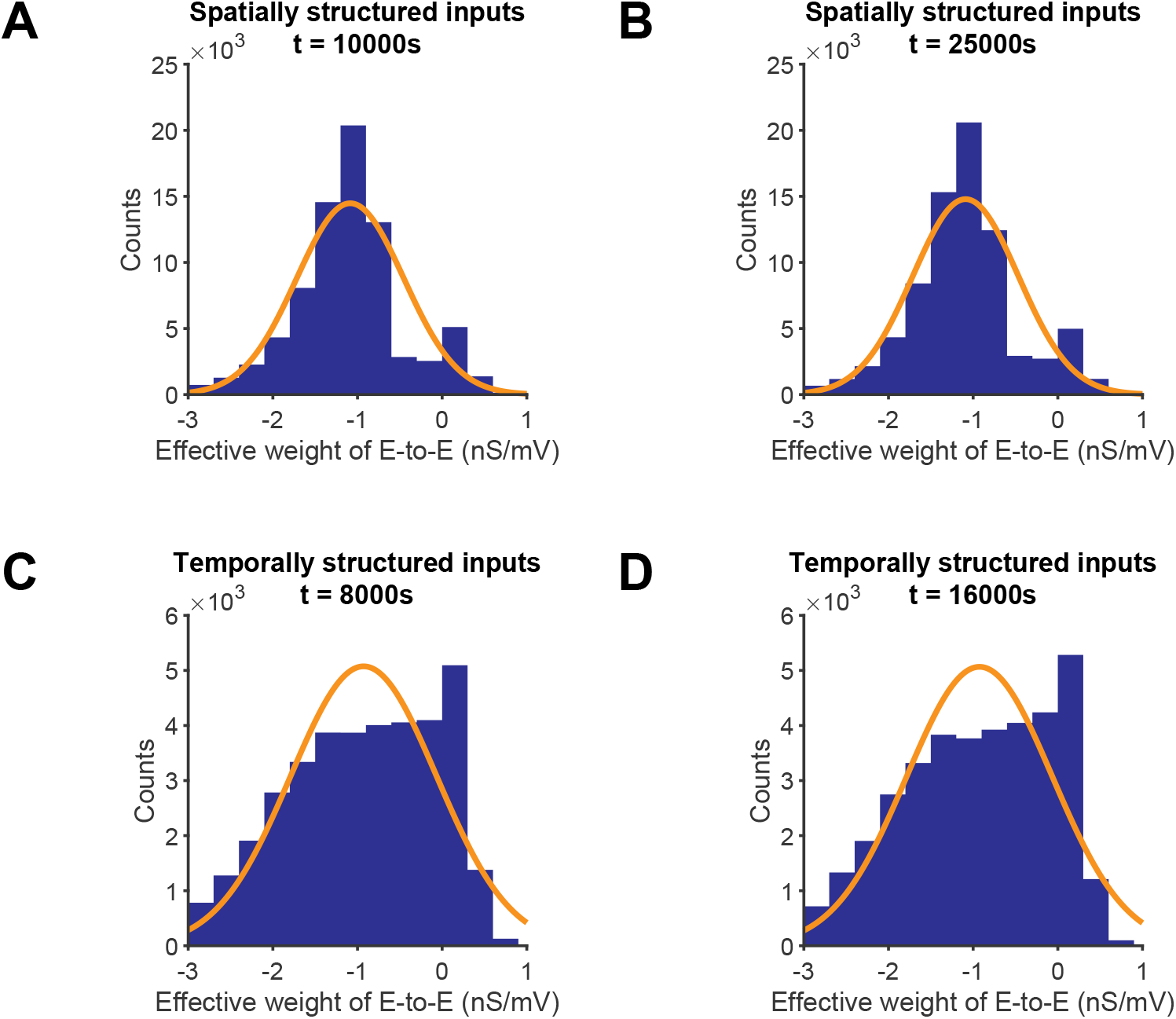
E-to-E distribution reached a steady state after training with structured inputs. **A**. The network develops a stable distribution which can be fit as lognormal when trained with spatially structured inputs. Left: *t* = 10, 000 s, right: *t* = 25, 000 s, orange curve shows the fit, *R*^2^ = 0.809 for the left and *R*^2^ = 0.821 for the right. **B**. The network develops a stable distribution which can be fit as lognormal when trained with temporally structured inputs. Left: *t* = 8, 000 s, right: *t* = 16, 000 s, orange curve shows the fit, *R*^2^ = 0.669 for the left and *R*^2^ = 0.669 for the right.

**Figure S12.**
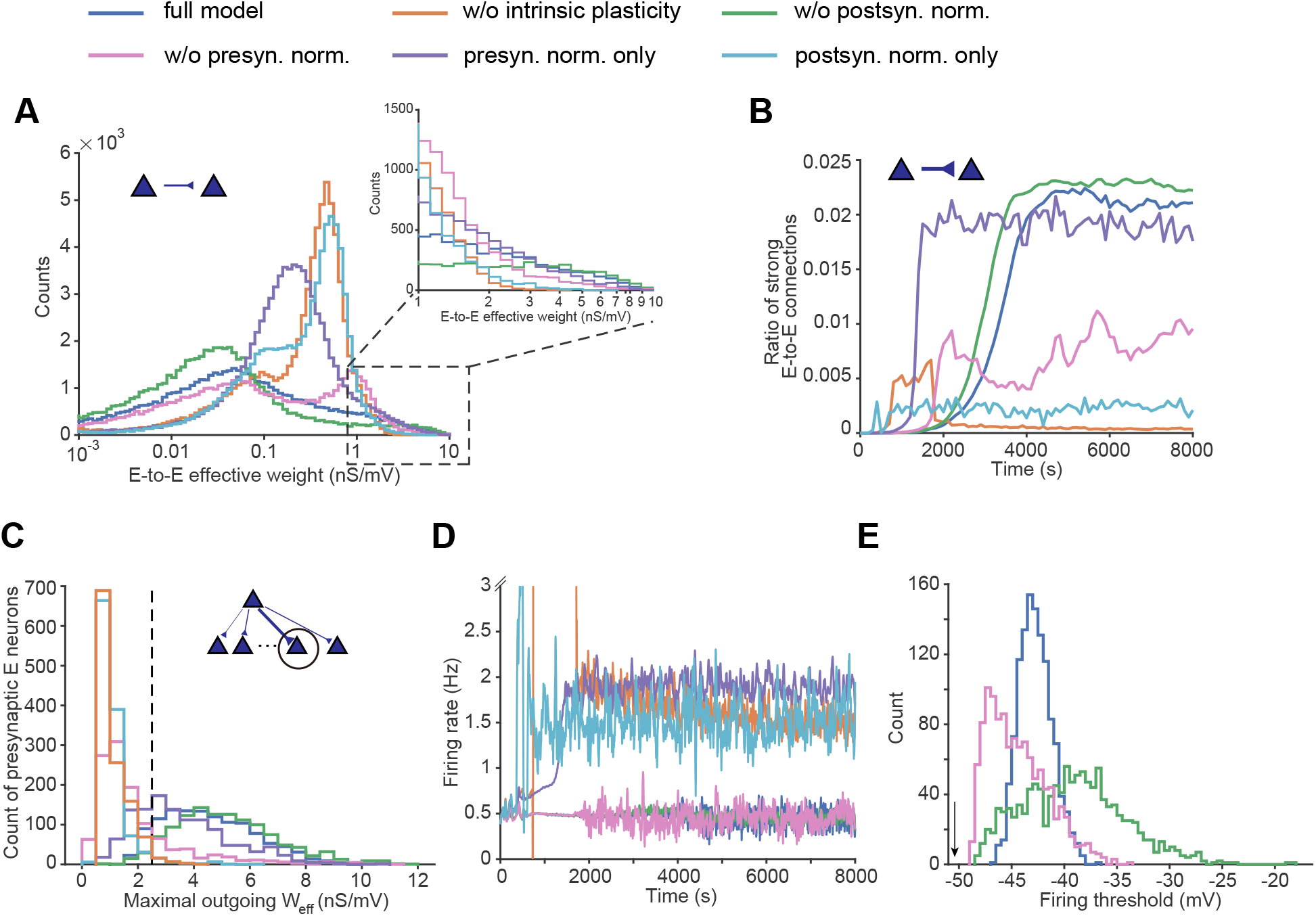
Alternative plasticity models can also support sequence-generating networks. **A**. Steady-state distribution of E-to-E synaptic weight under different combinations of plasticity rules (counterpart of Fig. 1C). Inset: Zoomed-in view of the distribution from 1 to 10 nS/mV. **B**. Ratio of strong E-to-E connections (*W*_eff_ *>* 2.5) over time (counterpart of Fig. 2B solid line). **C**. Histogram of the maximal outgoing *W*_eff_ across the 1,200 E neurons in the steady-state model network (counterpart of Fig. S2A, gray shade). The black dashed line indicates the minimal *W*_eff_ required to initiate a sequence (*W*_eff_ = 2.5). **D**. Traces of population firing rate of E neurons over time, smoothed using a third-order Savitzky-Golay filter with a 50-second time window. The y-axis is clipped at 3 Hz for visualization. **E**. Distribution of firing thresholds of E neurons in the steady-state network when intrinsic plasticity is active. Black arrow on the left indicates the initial firing threshold, which was identical for all excitatory neurons.

